# Identification and SAR optimization of FBXO22-mediated TEAD Targeted Glue™ degraders

**DOI:** 10.64898/2026.04.21.719895

**Authors:** Martin Fisher, Alice Fletcher, Callum J. Hamby, Jack T. N. Miles, Martin Ambler, Aleksandra Azevedo, Sara Bisetto, Colin T. R. Davies, Pauline Drouhin, Liliana Greger, Scott J. Hughes, Martin O’Rourke, Dominic D. G. Owens, Paula MacGregor, Wojciech J. Stec, Anna Tasegian, Martin Pass, Giles A. Brown, Louise K. Modis, Marta Carrara

## Abstract

TEAD transcription factors are emerging oncology targets due to their function as key effectors of the Hippo signalling pathway, which is frequently dysregulated in cancer. Here, we report the discovery and development of potent, deep, and rapid-acting TEAD Targeted Glue™ degraders that leverage an aldehyde-mediated degron mechanism. Our compounds demonstrate exceptional on-pathway selectivity profiles and exhibit the expected Hippo signalling modulation.

Mechanistically, we demonstrate that amine-based TEAD degrader scaffolds undergo extracellular conversion to the active aldehyde species, which mediate covalent engagement of FBXO22 C326, triggering TEAD proteasomal degradation. The degraders identified were able to be rationally and systematically optimised for both degradation potency and kinetics, achieving enhanced degradation profiles compared to previously reported FBXO22-targeting approaches, establishing design principles for this degrader class.

Our findings highlight strategies for the structure activity relationship (SAR) and rational optimization of aldehyde-mediated degrons to generate novel precision molecular glue degraders against TEAD, a high value oncology target.

## Introduction

Cells have evolved sophisticated protein quality control systems to recognise and eliminate damaged proteins through specific degradation signals. Targeted protein degradation (TPD) has emerged as a transformative therapeutic modality that harnesses the cellular ubiquitin-proteasome system to eliminate disease-causing proteins. The TPD field encompasses several strategies, the most prominent being proteolysis-targeting chimeras (PROTACs) and molecular glue degraders (MGDs), both of which induce proximity between target proteins and E3 ubiquitin ligases to promote proteasomal degradation ^1–6^.

Bivalent PROTAC degraders often exist in extended or beyond rule of 5 chemical space, meaning optimising their drug like properties is challenging. In contrast, MGDs typically only have affinity for the E3 ligase and lack a linker region which greatly improves their drug-like properties, such as oral bioavailability, while maintaining the ability to induce novel protein-protein interactions through small molecule stabilisation. However, despite their therapeutic potential, the development of MGDs has been faced with significant challenges, particularly their rational discovery and optimisation ^7,8^.

Cereblon (CRBN)-recruiting compounds represent the most prevalent MGD mode of action ^7,9–12^. Other E3 ligases are starting to be harnessed for MGD development, representing an expansion beyond CRBN-mediated degradation. Recent MGD advances have demonstrated the successful functional recruitment of diverse E3 ligases such as F-box only protein 22 (FBXO22) ^13–18^, DDB1 and CUL4 associated factor 16 (DCAF16) ^19–25^, DDB1 and CUL4 associated factor 15 (DCAF15) ^26^, F-box/WD repeat-containing protein 1A (β-TRCP) ^27^, von Hippel-Lindau disease tumour suppressor (VHL) ^28^, Kelch-like ECH-associated protein 1 (KEAP1) ^29^, and Seven in absentia homolog 1 (SIAH1) ^30,31^. This diversification of utilised E3 ligases holds considerable promise for expanding the druggable proteome and accessing protein targets that may be recalcitrant to CRBN-mediated degradation, ultimately broadening the therapeutic applications of MGDs. To enable the identification of Targeted Glue™ degraders, we deploy a distinctive target-first approach that fundamentally differs from conventional MGD discovery strategies. This approach enables the agnostic ‘glueing’ of the most suitable E3 ligase for a pre-selected, therapeutically valuable protein of interest (POI) through the incorporation of a minimal pharmacophore onto the POI-targeting ligand. In this work we describe the identification and optimisation of amine **1**, a novel Targeted Glue™ degrader of Transcriptional Enhanced Associate Domain (TEAD) that acts via FBXO22, the substrate recognition component of a SKP1-CUL1-F-box protein E3 ligase.

TEADs are a family of four transcription factors (TEAD1-4) that serve as key effectors of the Hippo signalling pathway. Under normal physiological conditions, the Hippo pathway maintains tissue architecture by preventing excessive cell growth through phosphorylation-mediated inactivation of the transcriptional co-activators Yes-Associated Protein (YAP) and Transcriptional Co-activator with PDZ-binding motif (TAZ). However, when the Hippo pathway is dysregulated - a frequent occurrence in cancer - YAP and TAZ translocate to the nucleus where they form transcriptionally active complexes with TEAD proteins ^32,33^. These TEAD-YAP/TAZ complexes drive the expression of genes involved in cell proliferation, survival, and stemness, including cellular communication network factor 1 (*CCN1*) and cellular communication network factor 2 (*CCN2*), ultimately promoting oncogenic transformation and tumour progression ^34^. TEAD proteins represent highly attractive therapeutic targets for solid tumours, particularly for mesothelioma and non-small cell lung cancer (NSCLC) where YAP/TAZ hyperactivation and subsequent TEAD-mediated transcription are frequently observed and correlate with poor prognosis and therapeutic resistance.

Perturbation of TEAD-YAP/TAZ transcriptional activity can be achieved either by small molecule inhibitors that bind the YAP-TEAD interface and subsequently directly disrupt their interaction - termed YAP-TEAD interface disruptors, or through allosteric disruption of this interaction by inhibition of palmitoylation of the lipid binding pocket. Examples of both approaches are now in clinical trials including YAP-TEAD interface disruptor IAG933 ^35,36^ and lipid binding-pocket inhibitor VT-3989 ^37^. CRBN/VHL-based PROTAC degraders of TEAD have also been reported based on both classes of TEAD inhibitors, however these have not advanced to the clinic ^38–40^.

Our mechanism of degradation, which is mediated via FBXO22 C326, is akin to that recently reported against nuclear receptor binding SET domain protein 2 (NSD2) ^17,18^, FK506-binding protein 12 (FKBP12) ^13^, SWI/SNF related, matrix associated, actin dependent regulator of chromatin, subfamily a, member 2/4 (SMARCA2/4) ^15,16^, bromodomain-containing protein 4 (BRD4) and cyclin-dependent kinase 12 (CDK12) ^14^. In contrast to those recent publications, we are also able to describe a route to rational optimisation of these compounds using standard medicinal chemistry techniques.

Crucially, (i) our degraders are small and glue-like with favourable physicochemical properties unlike TEAD PROTACs; (ii) our FBXO22-binding pharmacophore was able to be optimised rationally, despite its minimal size; and (iii) we demonstrate exceptional on-pathway modulation in cells. These findings establish aldehyde-mediated degradation as a powerful strategy for developing next-generation cancer therapeutics distinct from traditional E3 ligase recruitment strategies.

## Results

### Amine 1 potently and selectively mediates on-target TEAD degradation

Drug-like TEAD inhibitors fall into two classes: lipid binding-pocket inhibitors and YAP-TEAD interface disruptors. IAG933, a YAP-TEAD interface disruptor, has high binding affinity for all four TEAD paralogs with rapid and reversible binding kinetics ^35^. A related YAP-TEAD interface disruptor (Supp 1) was selected as the POI ligand. Amine **1** was identified as part of a broader hit finding campaign and was considered an attractive starting point for optimisation due to its potent and deep degradation activity (85% D_max_, 13 nM DC_50_) (Fig 1a) and the relatively small size of the putative degron tag. As a degrader, amine **1** had small-molecule-like physicochemical properties, particularly a lower molecular weight (655 Daltons (Da)) compared to TEAD PROTACs containing similar POI ligands linked to classical CRBN recruiting ligands (example shown in Supp 2) which are between 800-1000 Da ^39,40^.

**Figure 1:**
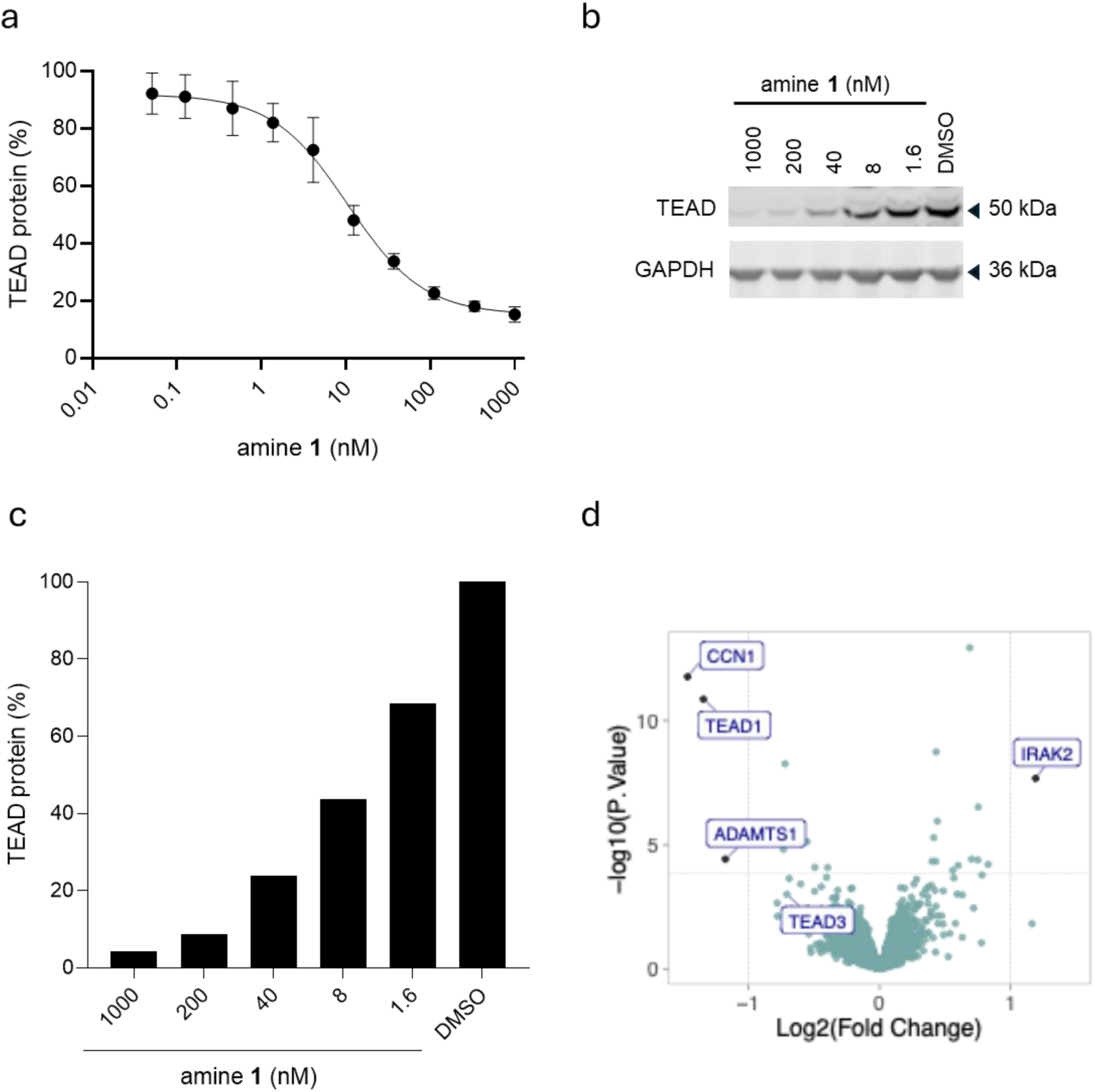
Potent and selective TEAD degradation by amine 1. **a**. TEAD protein levels in NCI-H226, determined by immunofluorescence, following 24 hours treatment with amine **1**. Data show the mean and SD from n = 3 biologically independent samples. **b-c**. Immunoblot and densitometry of pan-TEAD in NCI-H226 following 24 hours treatment with amine **1**. Data are from n = 1 biologically independent samples. **d**. Volcano plot displaying change in protein expression after 6 hours treatment with amine **1** vs DMSO in NCI-H226. Proteins were considered significantly decreased with a log_2_ fold change < −1 and adjusted p-value < 0.05.Hippo pathway associated proteins such as CCN1, ADAMTS1, TEAD1, TEAD3 are indicated in the plot.

Degradation of all TEAD isoforms present in NCI-H226 cells was subsequently validated by immunoblot analysis using a pan-TEAD antibody (Fig 1b, c). While RNA transcripts of all TEAD isoforms (TEAD1-4) have been detected in NCI-H226 cells (Cancer Dependency Map (DepMap)) ^41^, only TEAD1 and TEAD3 proteins were quantified in a global proteomics analysis of NCI-H226 cells (Fig 1d). Following 6 hours treatment with 500 nM amine **1,** out of the 7083 quantified proteins, the only significantly decreased proteins were TEAD1 (log_2_ fold change (FC) −1.34, p-adjusted 6.26e-9), CCN1 (log_2_ FC −1.46, p-adjusted 6.08e-9), and ADAMTS1 (log_2_ FC −1.17, p-adjusted 0.0195), known downstream target genes ^42,43^. TEAD3 was also downregulated, albeit not significantly at 6 hours (log_2_ FC −0.706, p-adjusted 0.163). Additionally, we ruled out non-specific bystander TEAD degradation by validating lack of TEAD degradation by amine **2** (Supp 3a, Supp Table 1), which contains a non-binding atropisomer of the POI ligand (amine **1** IC_50_ 1.8 nM, amine **2** IC_50_ > 4000 nM) (Supp 3b, Table 1, Supp Table 1). These data support on-target TEAD degradation by amine **1**.

**Table 1:**
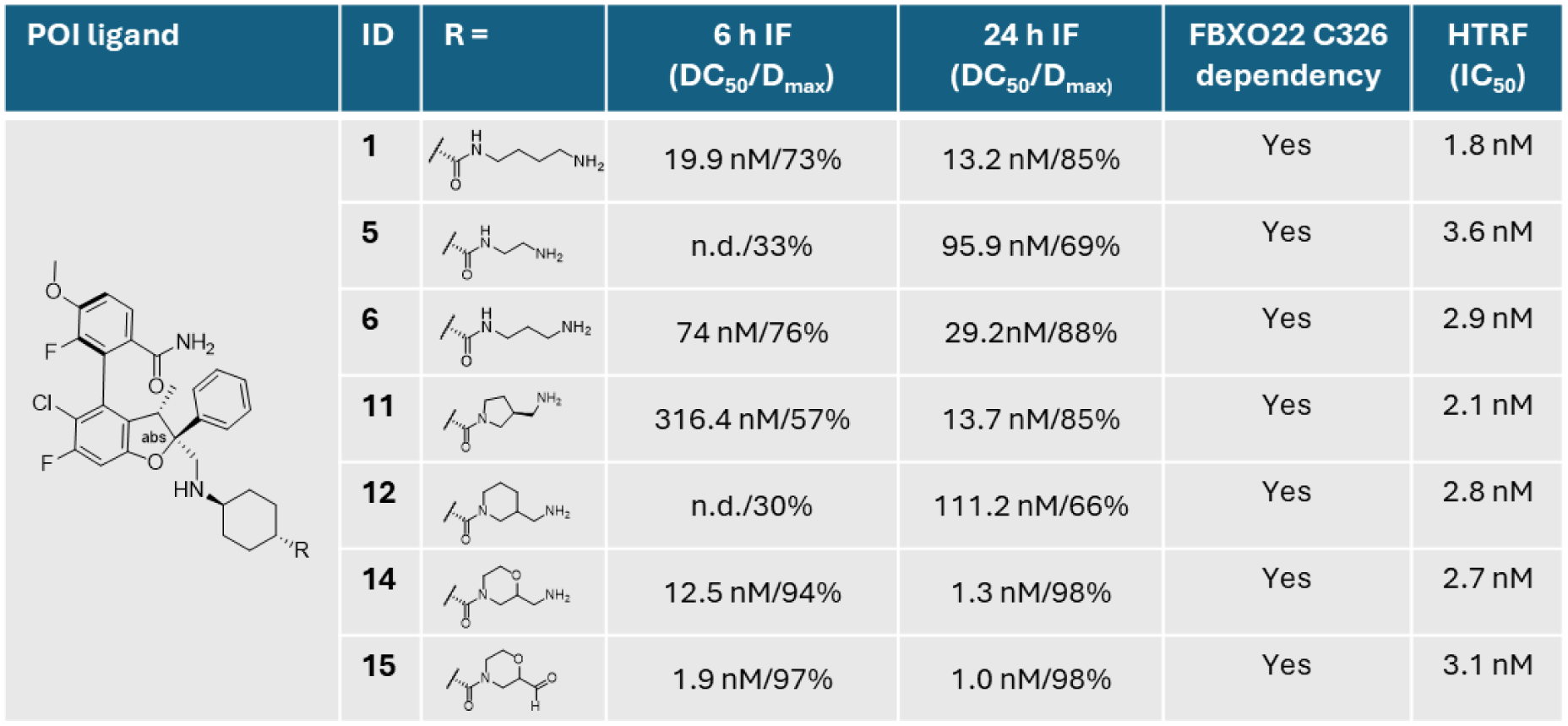
Tabulated data summary for TEAD degraders. Data summary for TEAD degraders amine **1,** amine **5,** amine **6,** amine **11,** amine **12,** amine **14** and aldehyde **15**. Activity at 6 hours and 24 hours in NCI-H226 as determined by immunofluorescence; FBXO22 dependency as determined by loss of activity in FBXO22 KO HeLa TEAD immunofluorescence; TEAD1 binding as measured via HTRF TEAD1-YAP displacement assay. IF data represents mean DC_50_ and D_max_ values for n≥2 biologically independent samples; FBXO22 C326 dependency was determined from n = 2 biologically independent samples; HTRF data is mean IC_50_ values of n≥3 biological replicates. Graph for amine **1** HTRF is shown in Supp 3b.

| POI ligand | ID | R = | 6 h IF<br>($DC_{50}/D_{max}$) | 24 h IF<br>($DC_{50}/D_{max}$) | FBXO22 C326<br>dependency | HTRF<br>($IC_{50}$) |
| --- | --- | --- | --- | --- | --- | --- |
|  | **1** |  | 19.9 nM/73% | 13.2 nM/85% | Yes | 1.8 nM |
|  | **5** |  | n.d./33% | 95.9 nM/69% | Yes | 3.6 nM |
|  | **6** |  | 74 nM/76% | 29.2nM/88% | Yes | 2.9 nM |
|  | **11** |  | 316.4 nM/57% | 13.7 nM/85% | Yes | 2.1 nM |
|  | **12** |  | n.d./30% | 111.2 nM/66% | Yes | 2.8 nM |
|  | **14** |  | 12.5 nM/94% | 1.3 nM/98% | Yes | 2.7 nM |
|  | **15** |  | 1.9 nM/97% | 1.0 nM/98% | Yes | 3.1 nM |

### Amine oxidation to aldehyde drives TEAD degradation

Considering the previously reported oxidation of primary amine groups in cellular environments, we explored the possibility that enzymatic conversion could yield a distinct active species ^13,15–17^. Co-treatment of 100 nM of amine **1** with aminoguanidine (AG) (a pan-amine oxidase inhibitor) prevented TEAD degradation in a dose-responsive manner, supporting a role for compound oxidation to aldehyde **3,** which represents the active species (Fig 2a,b, Supp 4a). Indeed, AG treatment had no effect on TEAD degradation by 100 nM aldehyde **3** (Fig 2b, Supp 4a). Chemical substitution of the amine in a manner which prevented oxidation, in this case methylation to form a secondary amine, also prevented degradation of TEAD (methyl amine **4**) (Supp Table 1). This conversion to the active species occurs extracellularly, as evidenced by the observation that amine **1**, unlike aldehyde **3**, loses its activity in serum-free conditions (Supp 4b).

**Figure 2:**
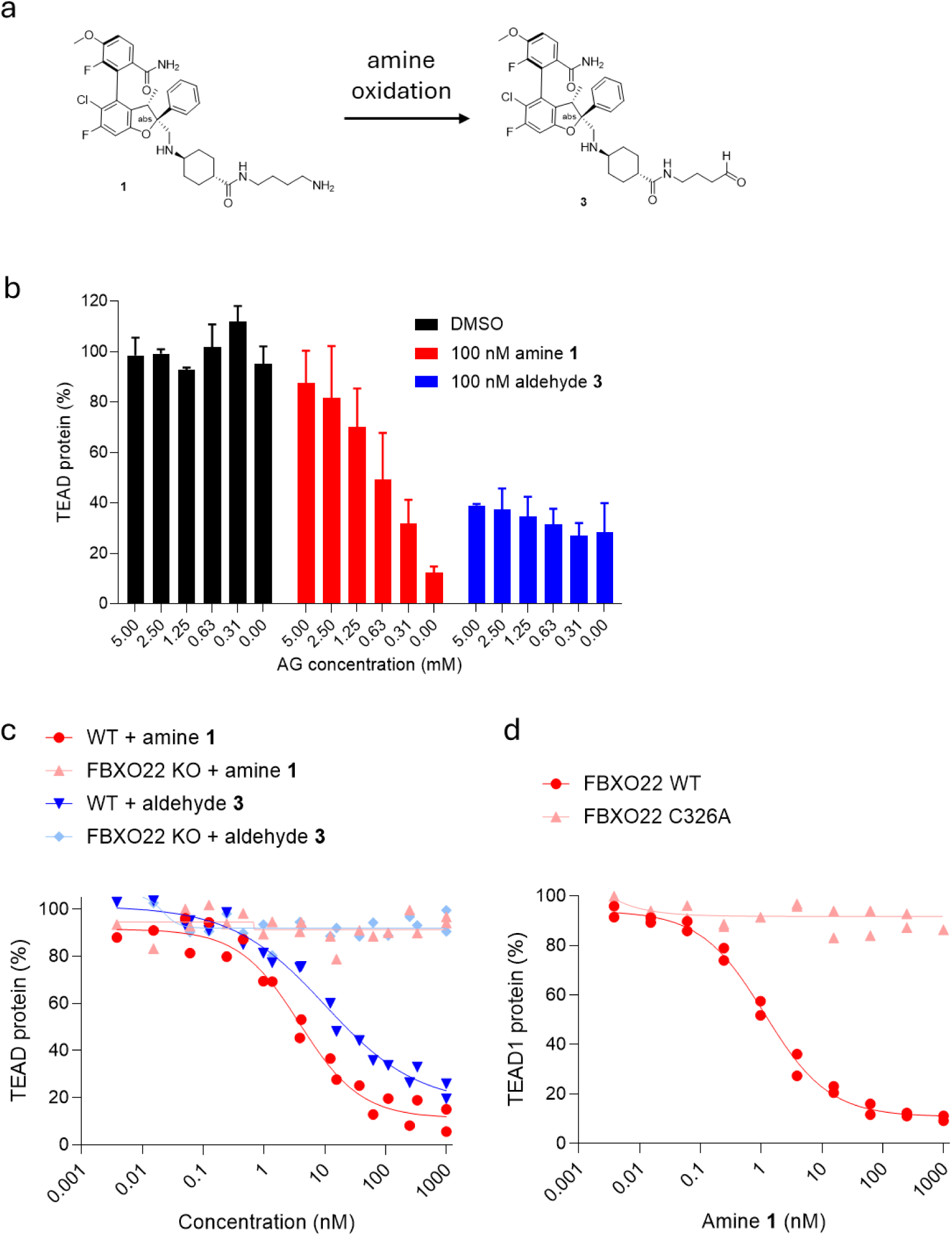
Amine 1 metabolism to its corresponding aldehyde 3 drives TEAD degradation via FBXO22. **a**. Chemical structure of amine **1** and its subsequent product aldehyde **3** following amine oxidation. **b**. Pan-TEAD protein levels in NCI-H226, determined by immunoblotting, following 24 hours treatment with 100 nM amine **1** or aldehyde **3** in the presence of AG. Data show the mean and SD from n = 2 biologically independent samples; one representative immunoblot shown in Supp 3a; additional blot shown in source data file. **c**. TEAD protein levels in parental HeLa WT or FBXO22 KO cell lines, determined by immunofluorescence, following 24 hours treatment with amine **1** or aldehyde **3**. Data show the individual data points from n = 2 biologically independent samples. **d**. TEAD protein levels in lentiviral HeLa FBXO22 KO + WT FBXO22 and HeLa FBXO22 KO + C326A FBXO22 cell lines, determined by immunofluorescence, following 24 hours treatment with amine **1**. Data are from n = 1 biologically independent samples.

### TEAD degradation by amine degraders is dependent on FBXO22 C326

The deep and potent degradation observed with amine **1** allowed it to be used as a tool compound to characterise the mechanism of action. We established that TEAD degradation was fully dependent on RING Cullin E3 activity, as demonstrated by the rescue of TEAD degradation upon cotreatment of amine **1** with MLN4924 - an inhibitor of the catalytic subunit of the NEDD8-activating enzyme (NAE-1) (Supp 5a).

Recent reports have shown that similar amine-based NSD2 ^17,18^, FKPB12 ^13^, SMARCA2/4 ^15,16^, BRD4 and CDK12^14^ degraders function via recruitment of FBXO22 E3 ligase. Indeed, by using isogenic wild-type (WT) and FBXO22 knock-out (KO) HeLa cells (Supp 5b) we were able to show dependency for amine **1** activity on FBXO22 E3 ligase (90% D_max_, 4.2 nM DC_50_ in HeLa WT; 5% D_max_ in HeLa FBXO22 KO) (Fig 2c). Similar results were obtained with aldehyde **3** (77% D_max_, 22.5 nM DC_50_ in HeLa WT; 5% D_max_ in HeLa FBXO22 KO) (Fig 2c). In contrast, a CRBN-recruiting TEAD PROTAC (PROTAC **16**) (structure shown Supp 2) retained activity in FBXO22 WT and KO (98% D_max_, 1.2 nM DC_50_ in HeLa WT; 97% D_max_, 2.2 nM DC_50_ in HeLa FBXO22 KO) (Supp 5c).

FBXO22 C326 has been shown to mediate targeted degradation by amine/aldehyde compounds ^13–18^. To assess the role of FBXO22 C326 in our degradation mechanism, we measured the activity of amine **1** in FBXO22 KO HeLa cells transduced to express WT or C326A FBXO22. Whilst amine **1** was inactive in FBXO22 KO cells, its ability to degrade TEAD was fully rescued upon re-expression of FBXO22 WT but not FBXO22 C326A, establishing an essential role for C326 in amine **1**-mediated TEAD degradation (Fig 2d). To ensure that the lack of degradation rescue by FBXO22 C326A was not due to lower expression levels or more general impaired function of the mutant protein, we validated that (i) expression levels were comparable amongst all constructs tested (Supp 5d), (ii) TEAD levels remained unchanged (Supp 5e) and (iii) that all proteins retained WT activity, as measured by their ability to degrade BACH1, a known FBXO22 substrate (Supp 5e) ^44,45^. Based on these results, and in line with previous reports, we propose a mechanism whereby a reactive aldehyde electrophile functions as a molecular damage-tag, effectively marking TEAD for recognition by the cellular protein quality control machinery, specifically C326 of FBXO22 E3 ligase.

### Rational optimisation of compounds for TEAD degrader activity

Identification of the active species and mechanism of action informed a design strategy to modify the position of the electrophile in relation to the surface of the TEAD protein to optimise activity. Initially, simple variations to linker length were explored. Both shortening (amines **5** and **6** (Table 1) and lengthening the alkyl chain (amines **7** and **8** (Supp Table 1)) from the amine to the amide were tolerated. The higher D_max_ of compounds **1** and **5** demonstrated the optimum linker length was between 3-4 carbon atoms from the amide (Table 1). Methylation of the linker was also tolerated but with significantly reduced activity, the more sterically bulky gem-dimethyl group was not tolerated and resulted in an inactive compound (amines **G** and **10**) (Supp Table 1).

Reducing linker flexibility was then investigated by cyclisation about the amide bond. Pyrrolidine rings such as cyclic amine **11** were favoured over the piperidine **12** and azetidine **13** but did not show a significant boost in activity compared to the acyclic amine **1** (Table 1, Supp Table 1). The amino morpholine **14**, however, was the most active compound identified with > 94% maximal degradation of TEAD achieved at both 6 hours (12.5 nM DC_50_), and 24 hours (1.3 nM DC_50_) (Table 1).

Despite significant changes in compound design and variation in the degradation profiles observed between compounds, the mechanism of degradation was consistent, with all compounds fully dependent on FBXO22. Significant changes in binding affinity to TEAD1 were not observed, with all compounds retaining low nM activity in a Homogeneous Time-Resolved Fluorescence (HTRF) assay (Table 1, Supp Table 1). This indicates, as would be expected, that variation in TEAD binding is not driving variation in degradation efficacy. The most likely explanation is that changes to the compound drive formation of different ternary complexes of differing productivity. Gratifyingly, degradation was sustained out to 72 hours (Fig 3a) suggesting that the electrophile is sufficiently stable to suppress TEAD degradation for longer than the measured half-life of around 24 hours (Supp 6a). In particular, aldehyde **15** demonstrated superior degradation kinetics, achieving > 50% TEAD degradation within 30 minutes of treatment and maintaining > 94% D_max_ at all time points beyond 2 hours (Fig 3a). These results represent both more rapid onset and greater extent of degradation compared to PROTAC **16** (Fig 3b). Together with reported NSD2 degraders such as UNC10088 ^17^, this constitutes the fastest reported degradation kinetics for FBXO22-mediated protein degradation in the literature to date, although our degraders reach D_max_ at 100x lower concentrations.

**Figure 3:**
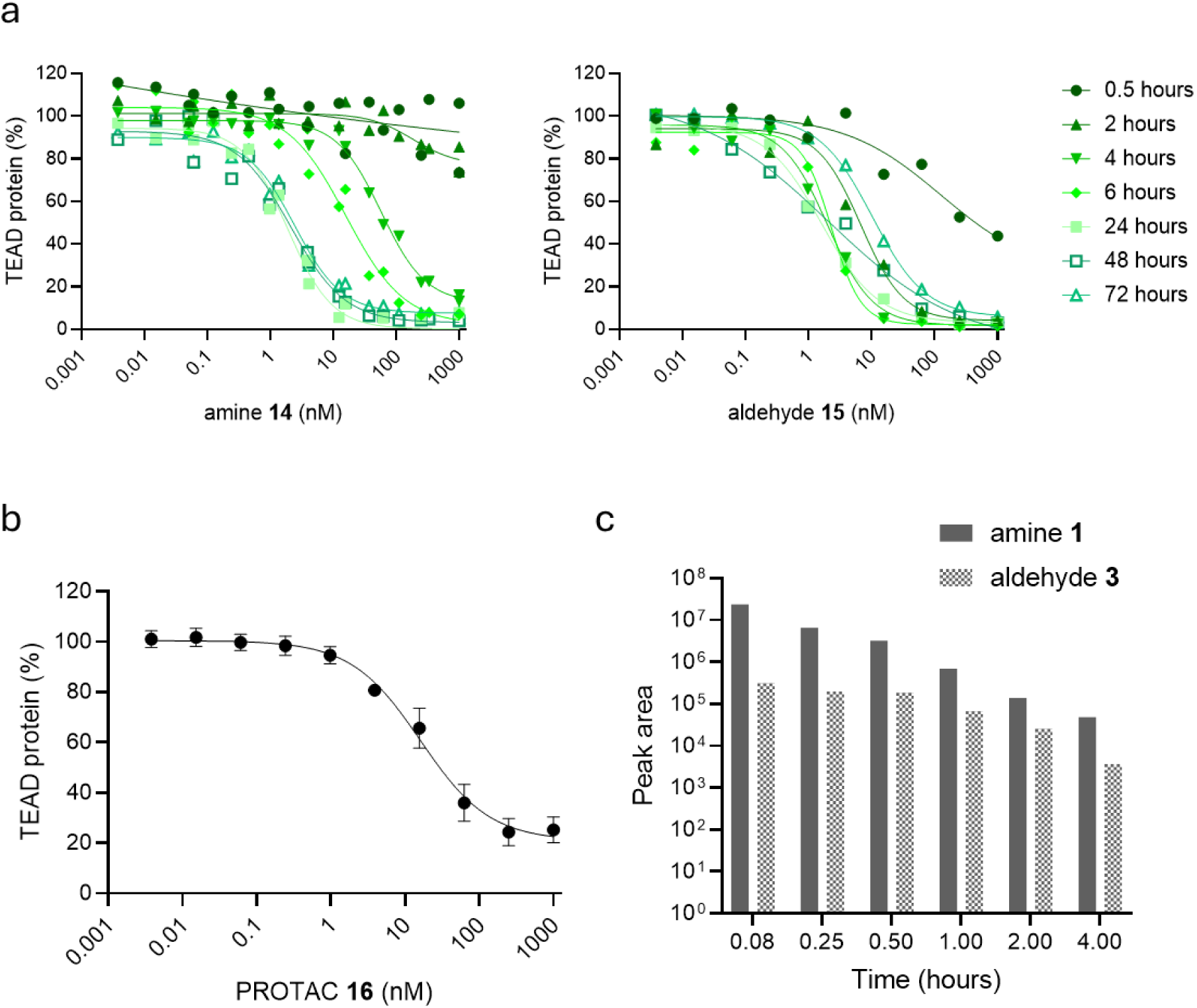
Rational compound design improves degrader activity. **a**. TEAD protein levels in NCI-H226, determined by immunofluorescence, following treatment with amine **14** and aldehyde **15** at a timepoints ranging from 0.5 to 72 hours. Data are from n = 2 and n = 1 biologically independent samples, respectively. **b.** TEAD protein levels in NCI-H226, determined by immunofluorescence, following 2 hours treatment with a dose response of PROTAC **16**. Data show the mean and SD from n = 3 biologically independent samples. **c.** In vivo formation of aldehyde **3** from amine **1**. Data shown the mean peak area as observed by LCMS plotted on a logarithmic scale vs time. Samples were collected from plasma of three independent CD-1 mice.

### Compounds can be designed to be orally bioavailable and the degradation active species forms *in vivo*

To further explore the scope of this mechanism we sought to determine the pharmacokinetic (PK) profile of amine **1** and whether the metabolic conversion to the aldehyde **3** was also observed *in vivo*. A portion of amine **1** was converted to the active species *in vivo* with aldehyde **3** identified in mouse plasma by comparison to an authentic sample (Fig 3c and Supp 7b). This suggests that with further optimisation, this type of amine could act as a degradation-inactive pro-drug. PK parameters showed modest clearance and moderate volume of distribution for amine **1** suggesting a reasonable starting point for optimisation (Supp 7a). However, amine **1** was not orally available most likely because it contains two basic nitrogens which would be protonated at biological pH. Reduction of the pKa of the pendent nitrogen, as in the difluoro amine **17** which is still an active TEAD degrader (Supp Table 1), showed 23% oral bioavailability in mouse (Supp 7c). This initial result demonstrates the ability to optimise amine-based degraders by simple modification of physicochemical properties in a similar manner to small molecule inhibitors or molecular glues.

### Optimised TEAD FBXO22-dependent degraders drive potent and deep on-target biology

Following these promising results, we sought to characterise the downstream effects of TEAD degradation by our FBXO22-dependent compounds. First, we validated the activity of the degraders in additional cell lines, including MSTO-211H mesothelioma cells (Fig 4a), NCI-H358 (Supp 8a) and PC9 NSCLC cells (Supp 8b). The compounds broadly retained both potency and depth of degradation across all cell lines tested (> 72% D_max_, < 80 nM DC_50_ for all compounds across cell lines), demonstrating that this mechanism is broadly applicable, and represents a viable degrader strategy for multiple cancer cell types. Secondly, we assessed the effect of the compounds on TEAD downstream pathways. Indeed, IAG933 and our TEAD degraders completely inhibited transcription of *CCN1* and *CCN2*, known TEAD-regulated genes, (< 2 nM IC_50_, > 90% I_max_) (Fig 4b, Supp 8c).

**Figure 4:**
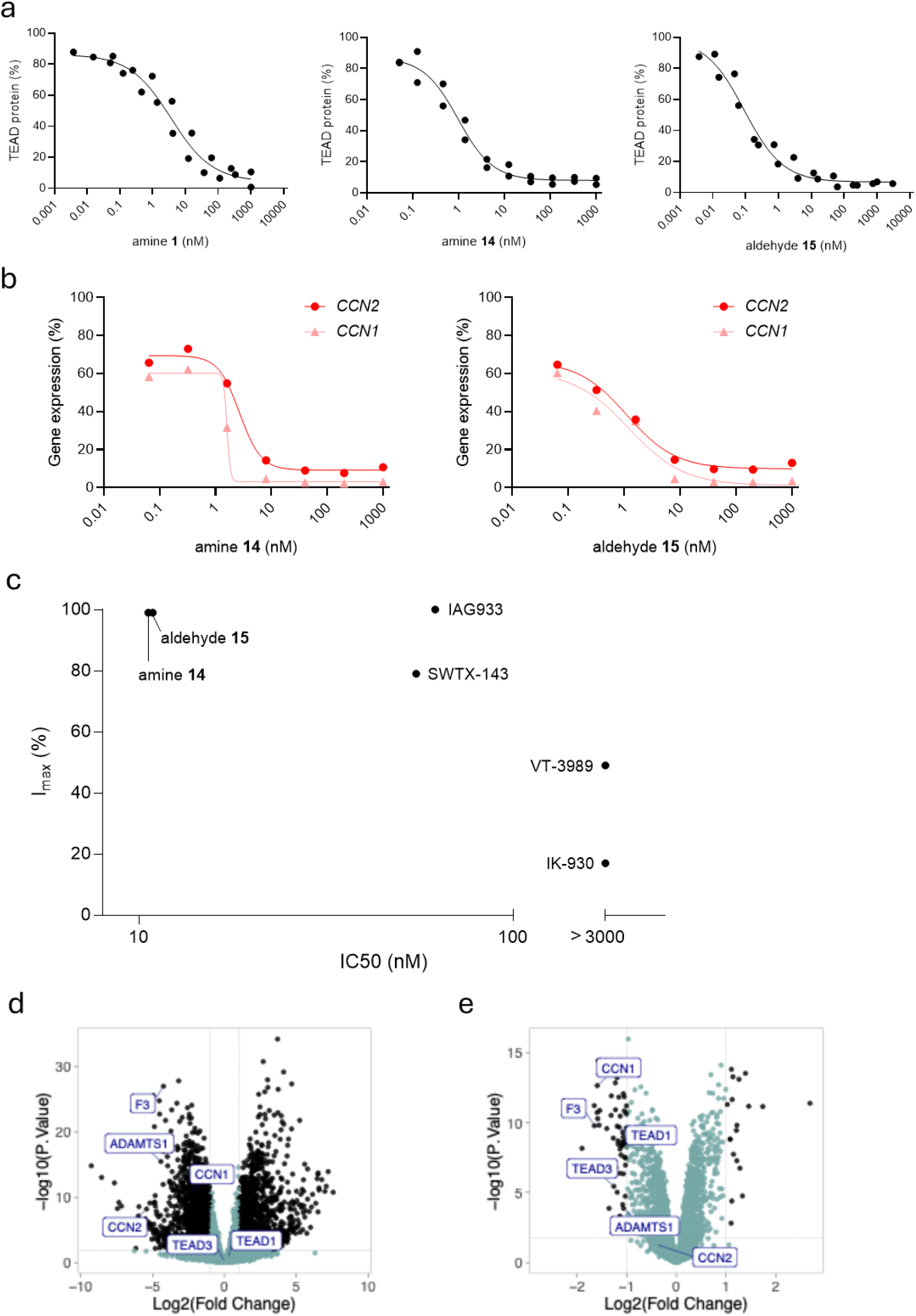
TEAD degraders drive efficacious on-target pathway modulation in mesothelioma cells. **a**. TEAD protein levels in MSTO-211H, determined by immunofluorescence, following 24 hours treatment with amine **1**, amine **14**, and aldehyde **15**. Data show the individual data points from n = 2 biologically independent samples. **b**. *CCN1* and *CCN2* gene expression levels measured by qPCR in MSTO-211H following 24 hours treatment with amine **1**, amine **14**, aldehyde **15**. Data are from n = 1 biologically independent sample. **c.** IC_50_ vs I_max_ of cell viability in MSTO-211H, determined by CellTiter-Glo, following 120 h treatment with a dose response of VT-3989, SWTX-143, IK-930, IAG933, amine **14** and aldehyde **15**. Curves are shown in Supp 10a. Data show the individual data points from n = 2 biologically independent samples and n = 1 biologically independent sample for SWTX-143. **d.** RNA-seq comparison between NCI-H226 cells treated for 24 hours with 500 nM of amine **1** or DMSO. Statistically significant genes were identified with p-adjusted < 0.05 and absolute log_2_ fold change > 1. Hippo pathway associated proteins TEAD1, TEAD3, CCN1, CCN2, F3, ADAMTS1 are labelled in the plot. **e.** Proteomics comparison between NCI-H226 cells treated for 24 hours with 500 nM of amine **1** compared to DMSO. Statistically significant proteins were identified with p-adjusted < 0.05 and absolute log_2_FC > 1. Hippo pathway associated proteins such as TEAD1, TEAD3, CCN1, CCN2, F3, ADAMTS1 are labelled in the plot.

Finally, MSTO-211H cells were utilised as a mesothelioma cell line with dysregulated Hippo pathway known to exhibit TEAD-dependency. IAG933, amine **14** and aldehyde **15** all showed > 95% maximal growth inhibition five days after treatment. Consistent with published data, this is higher than the maximal growth inhibition achieved by lipid binding-pocket inhibitors VT-3989, SWTX-143, and IK-930 (Fig 4c, Supp 9a). Similar depth of growth inhibition, and increased potency, was obtained with our optimised amine **14** and aldehyde **15** degraders. This cytotoxic effect was confirmed to be on-target, as demonstrated by the lack of effect of compounds on the viability of HepG2 Hippo pathway WT control cell line (Supp 9b).

### Selective TEAD degradation enables clean Hippo pathway modulation

To assess global transcriptional changes, RNA sequencing was performed 24 hours after treatment of cells with 500 nM (approximately 50-fold above 24 hours DC_50_) amine **1** (Fig. 4d). As expected, we observed downregulation of 1648 genes (log_2_ FC < 1, p-adjusted < 0.05), including *CCN1* and *CCN2*, confirming the compound’s impact on target gene expression. Gene set enrichment analysis (GSEA) confirmed on-target downregulation of known YAP/TAZ signature genes reported in the literature as well as known downstream pathways including E2F, G2M, and MYC targets ^32,46–48^. Amine **1** showed a similar transcriptional response to 24 hours treatment with 500 nM IAG933, with only four significantly differentially expressed genes between them (Supp 10a), further confirming the on-target transcriptional effect of amine **1**. To validate whether these transcriptional changes translated to the protein level, global proteomics experiments were conducted in parallel. These proteomic analyses corroborated the RNA-seq findings, demonstrating that transcriptional downregulation after amine **1** treatment resulted in corresponding decreases in protein abundance for known target genes including *CCN1*, *F3*, *ADAMTS1* (Fig 4e). As expected, TEAD1 and TEAD3, which are actively degraded upon amine **1** treatment, were only significantly modulated at the protein level, with no changes in transcript levels (Fig 4d,f). Again, protein changes after amine **1** treatment were similar to those obtained with 500 nM IAG933 treatment, except for TEAD1 and TEAD3 downregulation by our TEAD degrader only (Supp 10b). In addition, these findings are consistent with the transcriptomic and proteomic profiles observed following 24 hours treatment with 500 nM PROTAC **16** (Supp 10c,d), with no significant differences in protein levels between amine **1** and PROTAC **16**, (Supp 10e,f). Collectively, this indicates that the amine warhead does not confer additional off-target cellular perturbations beyond those attributable to the primary mechanism of action.

## Discussion

Here, TEAD degraders were identified through a target-first strategy that provides significant advantages for development of MGDs, enabling the rational selection of therapeutically valuable targets for degrader development. ^7,8^. This approach allows for the prioritisation of targets based on their therapeutic relevance rather than their compatibility with currently liganded E3 ligases, potentially accelerating the development of degraders against high-value targets that may not be amenable to traditional CRBN or VHL-based approaches. While CRBN-based degraders have shown remarkable clinical success ^49^, particularly in haematological malignancies, this narrow reliance on a single E3 ligase presents both opportunities and limitations for the field. The expansion of the functionally liganded E3 ligase toolbox represents a critical opportunity to broaden the therapeutic scope of TPD by accessing targets whose expression profiles, subcellular localisation, or ternary complex geometry may not be optimal for CRBN or VHL-mediated degradation. Indeed, the literature precedent for TEAD degraders, while limited, has focused exclusively on CRBN-based PROTACs ^38–40^. Recent work has demonstrated that CRBN PROTACs can achieve potent degradation of TEAD at nanomolar concentrations. However, the exclusive reliance on CRBN for TEAD degradation may limit the therapeutic potential of these approaches.

In this work we identify amine **1** which comprises a high-affinity YAP-TEAD interface ligand with a pendant primary amine. We demonstrate that this amine-based scaffold can be rationally developed into highly potent degraders and that other drug-like properties, such as oral bioavailability, can be rationally designed into these molecules. Mechanistically, we demonstrate that the POI ligand binds to TEAD with low nanomolar affinity, whilst the aldehyde electrophilic warhead is recognised by and forms a reversible covalent interaction with C326 of FBXO22. This interaction facilitates assembly of a productive ternary complex that promotes TEAD proteasomal degradation.

Compound optimisation was facilitated by the consistent FBXO22 C326-mediated mechanism of action which led to clear structure activity relationships which could then be readily exploited. For example, substitution alpha to the pendent amine was not tolerated. Through systematic optimisation of amine **1** we established clear principles governing optimal linker length and demonstrated that linker rigidification can further modulate degrader activity enabling efficient optimisation from the initial hit. Importantly, the small size of the previously unreported morpholine-amide FBXO22-recruiting group allows this highly potent degrader to maintain attractive physicochemical properties despite the relatively large and complex POI ligand. This could potentially extend the aldehyde-degron approach to a wider variety of POI ligands, whilst maintaining small molecule-like physiochemical properties, unlike bifunctional degrader-based molecular designs.

Optimised aldehyde **15** demonstrates achieves deep (> 95% D_max_) and potent (< 10 nM DC_50_) degradation within 2 hours an exceptional degradation profile for any MGD or PROTAC and an exciting proof-of-concept for an FBXO22 mediated degrader. Using our optimised TEAD degraders, we show that our compounds achieve superior on-target Hippo pathway gene modulation compared to clinical-stage TEAD inhibitors, translating to more potent killing of mesothelioma cells *in vitro*. We further validated the expected on-target biology of TEAD degradation by measuring gene modulation and on-target cell killing in relevant cancer cell lines.

This work presents the extensive SAR development of FBXO22 C326-recruiting aldehyde warheads to generate potent, deep and rapid TEAD degraders with therapeutic potential in Hippo mutated cancers. Future studies will determine whether comparable activity can be retained upon replacement of the aldehyde functionality with alternative FBXO22-recruiting electrophilic warheads.

## Acknowledgements

The authors would like to thank John Goodall, James T. Lynch, Nikolai Makukhin and Andrea Testa for valuable discussions; Dr Kate Heesom (Director of the Bristol University Proteomic Facility, UK) and Viva Biotech for their technical assistance with this work; Krishna Reddy Annadi, Manas Pattanayak, Mohana Rao Vippila and Ramesh G.B Khanna and the team of scientists at Sai life Sciences Hyderabad for their work synthesising the molecules discussed in this work.

## Methods

### Cell culture and cell engineering

The human NCI-H226 (#CRL-5826) and MSTO-211H (#CRL-2081) mesothelioma cell lines, human NCI-H358 (#CRL-5807) NSCLC cell line and human HepG2 hepatocellular cell line (#HB-8065) were obtained from the ATCC. The human PC-9 (#90071810) NSCLC cell line was obtained from the ECACC. All cell lines were routinely passaged and seeded in RPMI medium containing glutaMAX (Gibco, #12027599) supplemented with 10% FBS (Gibco, #17479633) with the exception of the HepG2 which were cultured in DMEM containing glutaMAX (Gibco, #12077549) supplemented with 10% FBS (Gibco, #17479633). For the experiment completed without serum, medium was not supplemented with serum. For immunofluorescence, cells were seeded in phenol red-free RPMI (Gibco, #11835030) containing 10% FBS.

The human HeLa cervical carcinoma cell line was edited for FBXO22 KO, alongside a mock transfected control (WT) via CRISPR (Synthego/EditCo) and single clone populations were obtained. HeLa cell lines were routinely passaged and seeded in DMEM medium containing glutaMAX (Gibco, # 12077549) supplemented with 10% FBS (Gibco, #17479633). For immunofluorescence, cells were seeded in phenol red-free DMEM (Gibco, #P04-03591) containing 10% FBS.

Lentiviral cell lines were generated by transducing HeLa FBXO22 KO cell lines with FBXO22 WT or FBXO22 C326A (VectorBuilder), polyclonal populations were obtained.

The identity of cell lines was confirmed by STR by the vendor at purchase. All cell lines underwent routine mycoplasma testing using the MycoAlert Mycoplasma Detection Kit (Lonza, #LT07-318). All cell lines were maintained at 37°C and 5% CO_2_ in a humidified environment.

### Cell treatments

Stock solutions of compounds were prepared at a concentration of 10 mM in DMSO and stored at −20°C. Stock solutions were pre-diluted in DMSO to 1000-fold the desired top start concentration. Dilution series were prepared from stock solutions, as required.

MLN4924 (TOCRIS, #6499, dissolved to 10 mM in DMSO) stock was used for final concentration 3 µM; cycloheximide (ThermoFisher Scientific, #J66004.XF, diluted to 10 mg/ml in media) was used for final concentration 10 µg/ml; AG (Sigma-Aldrich, #396494) was solubilised to 15 mM in complete medium, made fresh for each experiment.

For high throughput assays, compounds were dispensed directly into cell assay plates using the FlexDrop iQ Non-contact Dispenser (Revvity). For 6-well plates, compound was manually pipetted. The final concentration of DMSO was 0.1%, or 0.2% for co-treatment with MLN4924.

### Immunofluorescence

We utilised an immunofluorescence-based assay to quantify TEAD1 to enable degrader identification in cells. Pan-TEAD antibodies were not suitable for use in this assay.

Cells were seeded in phenol red-free medium in 384-well black, optically clear PhenoPlates (Revvity, #6057300) and incubated overnight at 37°C and 5% CO_2_. The next day, cells were treated with compounds including any additional treatment (i.e. 3 µM MLN4924; 1 hour pre-incubation), as required. Treated cells were incubated for the defined treatment time at 37°C and 5% CO_2_.

Cells were then fixed with 16% methanol-free paraformaldehyde (ThermoFisher Scientific, #2890 [v/v 3.7%] and blocked prior to incubation with primary antibodies at 4°C overnight then secondary antibodies, including Hoechst to stain nuclei, at 37°C for 1 hour. All antibodies are described in Table 2. To measure protein levels, plates were imaged with the Operetta CLS High-Content Analysis System (Revvity).

**Table 2.**
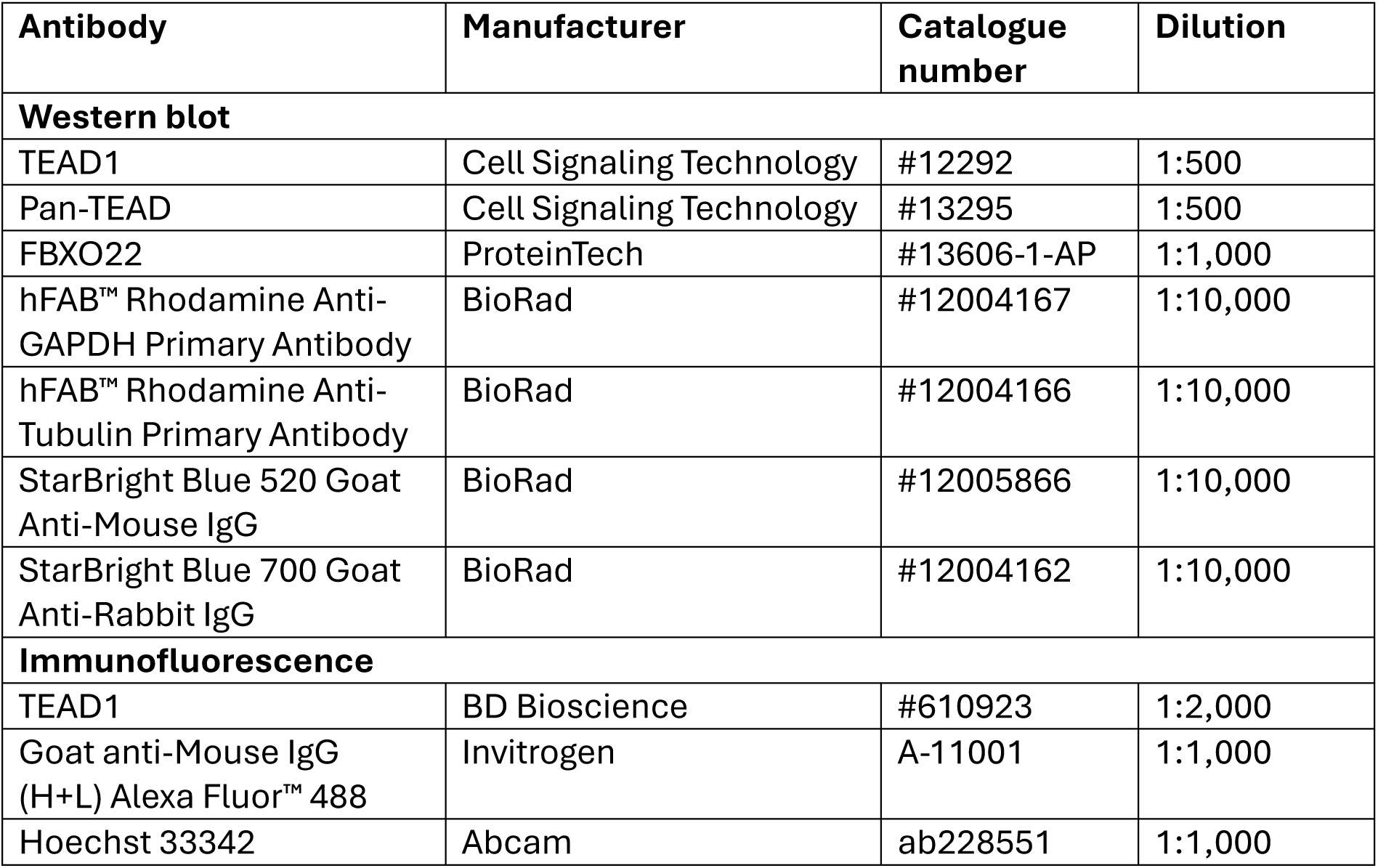
Immunofluorescence and immunoblot antibodies.

| Antibody | Manufacturer | Catalogue number | Dilution |
| --- | --- | --- | --- |
| **Western blot** |  |  |  |
| TEAD1 | Cell Signaling Technology | #12292 | 1:500 |
| Pan-TEAD | Cell Signaling Technology | #13295 | 1:500 |
| FBXO22 | ProteinTech | #13606-1-AP | 1:1,000 |
| hFAB™ Rhodamine Anti-GAPDH Primary Antibody | BioRad | #12004167 | 1:10,000 |
| hFAB™ Rhodamine Anti-Tubulin Primary Antibody | BioRad | #12004166 | 1:10,000 |
| StarBright Blue 520 Goat Anti-Mouse IgG | BioRad | #12005866 | 1:10,000 |
| StarBright Blue 700 Goat Anti-Rabbit IgG | BioRad | #12004162 | 1:10,000 |
| **Immunofluorescence** |  |  |  |
| TEAD1 | BD Bioscience | #610923 | 1:2,000 |
| Goat anti-Mouse IgG (H+L) Alexa Fluor™ 488 | Invitrogen | A-11001 | 1:1,000 |
| Hoechst 33342 | Abcam | ab228551 | 1:1,000 |

Image analysis was based on cellular fluorescent TEAD1 signal in individual cell nuclei, as identified by the Hoechst stain. Quantification was completed using Harmony imaging and analysis software (Revvity, V.5.2.), and data were normalised to DMSO and the no primary antibody control (background) to give “% protein”. For dose responses, absolute DC_50_ values were determined using a standard four parameter curve fitting, and D_max_ was considered as 100 – Y_min_.

### Immunoblotting

Cells were seeded in 6-well plates (Greiner BioOne, #657160) and incubated overnight at 37°C and 5% CO_2_. The next day, cells were treated with compounds including any additional treatment (i.e. AG; co-treatment with AMPH compounds), as required. Treated cells were incubated for the defined treatment time at 37°C and 5% CO_2_. Cells were lysed in RIPA buffer (ThermoScientific, #10017003) containing protease/phosphatase inhibitor (ThermoScientific, #15662249). Protein lysates were quantified via BCA (Fisher Scientific, #10056623) and prepared in 4x Laemmli SDS sample buffer (Fisher Scientific, #15492859) with DTT prior to denaturation at 95°C.

Proteins were resolved by SDS-PAGE then transferred onto 0.45 µM nitrocellulose membrane and blocked. Membranes were probed with primary antibodies outlined at 4°C overnight then secondary antibodies at room temperature for 1 hour. All antibodies are described in Table 2. Proteins of interest were detected with the ChemiDoc Imaging System (BIORAD). Quantification was completed using Image Lab (BIORAD, V.6.1.0), data were normalised to the loading control and calculated relative to DMSO.

### Quantitative PCR assay

Cells were seeded in 96-well plates (ThermoFisher Scientific, #167425) and incubated overnight at 37°C and 5% CO_2_. The next day, cells were treated with compounds and incubated for 24 hours at 37°C and 5% CO_2_.

Cells were washed with PBS then lysed with lysis buffer from the TaqMan™ Fast Advanced Cells-to-CT™ Kit (ThermoFisher Scientific, #A35374) according to the manufacturer’s protocol. 10 µL lysate was then added to 40 µL retro transcription master mix, and incubated on a thermocycler (SimpliAmp, ThermoFisher Scientific) following the manufacturer’s recommendation. Duplex reactions for quantitative PCR using 3 µL cDNA, TaqMan™ Fast Advanced Master Mix and TaqMan™ Gene Expression Assays – defined in Table 3 – were prepared then run on the real-time PCR instrument (QuantStudio 7Flex, ThermoFisher Scientific).

**Table 3.** TaqMan™ Gene Expression Assay.

| Gene, species | Manufacturer | Assay ID |
| --- | --- | --- |
| CTGF, human | Life Technologies | Hs00170014_m1 |
| Cyr61, human | Life Technologies | Hs00155479_m1 |
| GAPDH, human | Life Technologies | Hs99999905_m1 |
| RPLP0, human | Life Technologies | Hs99999902_m1 |

Gene expression was quantified using the 2^ΔΔCt^ method. Data were normalized to *RPLP0* and *GAPDH* housekeeping genes and values were averaged. Each data point represents the average for duplicate samples. Absolute EC_50_ values were determined using a standard four parameter curve fitting, and E_max_ was considered as 100 – Y_min_.

### Cell viability assay

Cells were seeded in 384-well plates (Greiner, #781080) and incubated overnight at 37°C and 5% CO_2_. The next day, cells were treated with compounds and incubated for 120 hours at 37°C and 5% CO_2_. Cell viability was assessed using CellTiter-Glo® 2.0 Cell Viability Assay (Promega, #G9243). Plates were equilibrated to room temperature prior to addition of CellTiter-Glo reagent in a 1:2 ratio to cell medium. Plates were placed in a shaker for 3 minutes then incubated at room temperature for a further 10 minutes to stabilize the luminescent signal. Luminescence was detected using the PHERAStar FSX (BMG LABTECH). Data were normalised to DMSO and 10 µM Staurosporine (background) to give “% viability”. For dose responses, absolute IC_50_ values were determined using a standard four parameter curve fitting, and I_max_ was considered as 100 – Y_min_.

### Global proteomics/RNA-seq cell treatments

NCI-H226 cells were seeded the day before at 1.5×10^6 for 6 hours and 24 hours samples and 1.0×10^6 cells for 48 hour samples in T75 flasks. The following day, cells were treated with 500 µM of indicated compounds. At the endpoint, cells were lifted with accutase, washed in PBS and divided into two samples (proteomics + RNA-seq) and pelleted at 300xg for 5 minutes. Proteomic samples were resuspended in 200 uL of RIPA, 2 uL of Benzonase and protease inhibitors (Pierce A32961). RNA-seq samples were resuspended in 350 µL of RLT plus buffer (Qiagen, 1053393). Samples were collected in biological triplicate.

### FASP-based Tryptic Digestion

Protein quantification using a Bradford assay was performed on lysates and 246 µg of each sample was used for FASP as follows: Dithiothreitol was added to a final concentration of 83.3 mM, incubated at 99°C for 5 minutes, then cooled to room temperature. VIVACON 500 filter units, 30,000 MWCO (Sartorius Stedim, Biotech GmbH, 37079 Goettingen, Germany) were washed with 8 M urea in 100 mM Tris/HCl pH 8.5, centrifuged at 14,000 × g, 20°C for 15 minutes. Samples were loaded in 50 µL aliquots plus 200 µL 8 M urea, centrifuged at 14,000 × g, 20°C for 15 minutes. The filter units were washed with 8 M urea, centrifuged at 14,000 × g, 20°C for 15 minutes. 100 μL 50 mM Iodoacetamide in 8 M urea was added to the filter units and incubated for 30 minutes in the dark. Filter units were centrifuged at 14,000 × g, 20°C for 10 minutes, then washed with 100 μL 8 M urea, centrifuged at 14,000 × g, 20°C for 15 minutes. This wash step was repeated two times, for a total of three washes. Filters were further washed with 100 μL 50 mM Triethylammonium Bicarbonate Buffer pH 8.5 (TEAB, (Thermo Fisher Scientific)), centrifuged at 14,000 × g, 20°C for 10 minutes. This wash step was repeated two times, for a total of three washes. Filter units were transferred to new tubes and 1.25% (w/w) trypsin (Pierce MS grade, Thermo Fisher Scientific, Loughborough, LE11 5RG, UK) in 60 µL 50 mM TEAB was added. The Filter units were sealed with Parafilm and digestion performed overnight at 37°C with shaking. Following overnight digestion, the filter units were centrifuged at 14,000 × g, 20°C for 20 minutes. and the flow through containing the peptides was retained. The filter units were washed with 40 μL 50 mM TEAB, centrifuged at 14,000 × g, 20°C for 10 minutes and then 50 μL 0.5M NaCl, centrifuged at 14,000 × g, 20°C for 20 minutes and these washes were added to the flow through. Peptide samples were desalted and cleaned up using Sep-Pak cartridges according to the manufacturer’s instructions (Waters, Milford, Massachusetts, USA). Eluate from the Sep-Pak cartridge was evaporated to dryness and resuspended in 100 µL 50 mM TEAB and 10 µL used for a peptide assay. All chemicals from Merck Life Science UK Limited, Dorset, SP8 4XT unless otherwise stated.

### TMT Labelling, High pH reversed-phase chromatography

For expression proteomics samples, approximately 60 µg of each sample was labelled with Tandem Mass Tag (TMTpro) 18-plex reagents according to the manufacturer’s protocol (Thermo Fisher Scientific, Loughborough, LE11 5RG, UK). The labelled samples were split across two experimental runs, each containing 9-plex experiments. The labelling scheme is as follows:

In Experiment 1, the samples were labelled as follows: TMT-126 = DMSO 6 hours Repeat 1, TMT-127C = DMSO 6 hours Repeat 2, TMT-128C = DMSO 24 hours Repeat 1, TMT-129C = DMSO 24 hours Repeat 2, TMT-130C = 500 nM amine **1** 6 hours Repeat 1, TMT-131C = 500 nM amine **1** 6 hours Repeat 2, TMT-132C = 500 nM amine **1** 24 hours Repeat 1, TMT-133C = 500 nM PROTAC **16** 24 hours Repeat 1, and TMT-134C = 500 nM IAG933 24 hours Repeat 1.

In Experiment 2, the samples were labelled as follows: TMT-126 = DMSO 6 hours Repeat 3, TMT-127C = DMSO 24 hours Repeat 3, TMT-127N = 500 nM PROTAC **16** 24 hours Repeat 2, TMT-128N = 500 nM PROTAC **16** 24 hours Repeat 3, TMT-130C = 500 nM IAG933 24 hours Repeat 2, TMT-131C = 500 nM IAG933 24 hours Repeat 3, TMT-131N = 500 nM amine **1** 24 hours Repeat 2, TMT-132N = 500 nM amine **1** 24 hours Repeat 3, and TMT-133C = 500 nM amine **1** 6 hours Repeat 3.

### Offline HpRP fractionation

For expression proteomics, samples were combined to a total of 100 µg each TMT plex and desalted using a SepPak cartridge according to the manufacturer’s instructions (Waters, Milford, Massachusetts, USA). Eluate from the SepPak cartridge was evaporated to dryness and resuspended in buffer A (20 mM ammonium hydroxide, pH 10) prior to fractionation by high pH reversed-phase chromatography using an Ultimate 3000 liquid chromatography system (Thermo Fisher Scientific). In brief, the sample was loaded onto an XBridge BEH C18 Column (130 Å, 3.5 µm, 2.1 mm X 150 mm, Waters, UK) in buffer A, and peptides were eluted with an increasing gradient of buffer B (20 mM Ammonium Hydroxide in acetonitrile, pH 10) from 0-95% over 60 minutes. The resulting fractions (20 in total) were evaporated to dryness and resuspended in 1% formic acid prior to analysis by nano-LC MSMS using an Orbitrap Fusion Lumos mass spectrometer (Thermo Scientific).

### Nano-LC MS3

High pH RP fractions were further fractionated using an Ultimate 3000 nano-LC system in line with an Orbitrap Fusion Lumos mass spectrometer (Thermo Scientific). In brief, peptides in 1% (vol/vol) formic acid were injected onto an Acclaim PepMap C18 nano-trap column (Thermo Scientific). After washing with 0.5% (vol/vol) acetonitrile 0.1% (vol/vol) formic acid peptides were resolved on a 250 mm × 75 μm Acclaim PepMap C18 reverse phase analytical column (Thermo Scientific) over a 150 minutes organic gradient, using 7 gradient segments in solvent B (1-6% for 1 minute, 6-15% for 58 minute, 15-32% for 58 minutes, 32-40% for 5 minutes, 40-90% for 1 minute, held at 90% for 6 minutes and then reduced to 1% over 1 minute) with a flow rate of 300 nL minute^−1^. Solvent A was 0.1% formic acid and Solvent B was aqueous 80% acetonitrile in 0.1% formic acid. Peptides were ionized by nano-electrospray ionization at 2.0kV using a stainless-steel emitter with an internal diameter of 30 μm (Thermo Scientific) and a capillary temperature of 300°C. For expression proteomics, 20 samples were collected for SPS-MS3 analysis.

All spectra were acquired using an Orbitrap Fusion Lumos mass spectrometer controlled by Xcalibur 3.0 software (Thermo Scientific) and operated in data-dependent acquisition mode using an SPS-MS3 workflow. FTMS1 spectra were collected at a resolution of 120,000, with an automatic gain control (AGC) target of 200,000 and a max injection time of 50 ms. Precursors were filtered with an intensity threshold of 5,000, according to charge state (to include charge states 2-7) and with monoisotopic peak determination set to Peptide. Previously interrogated precursors were excluded using a dynamic window (60s +/−10 ppm). The MS2 precursors were isolated with a quadrupole isolation window of 0.7 m/z. ITMS2 spectra were collected with an AGC target of 10,000, max injection time of 70 ms and CID collision energy of 35%.

For FTMS3 analysis, the Orbitrap was operated at 50,000 resolution with an AGC target of 50,000 and a max injection time of 105 ms. Precursors were fragmented by high energy collision dissociation (HCD) at a normalised collision energy of 60% to ensure maximal TMT reporter ion yield. Synchronous Precursor Selection (SPS) was enabled to include up to 10 MS2 fragment ions in the FTMS3 scan.

### Proteomics bioinformatics analysis

Raw proteomics data files were converted to mzML format using msconvert proteowizard (version 3.0.22167) before processing with FragPipe (version 21.1) and MSFragger (version 4.0) ^50^. Database search was performed against Human Uniprot reviewed reference proteome (2023_05, downloaded 5^th^ January 2024). Fixed modifications included were carbamidomethylation of cysteine (+57.02146 Da) and TMTpro labelling of Lysine (+304.20715 Da). Variable modifications included were Oxidation of methionine (+15.9949 Da), N-terminal acetylation (+42.0106 Da), and N-terminal TMTpro labelling (+304.20715 Da). False discovery rate control was set to 1% at the PSM, peptide, and protein level. Total channel normalization was performed to account for channel loading differences and virtual reference normalization was performed by using internal reference scaling to the average intensity in each plex to combine the two plexes. Unique peptides were used for the quantification of detected TEAD proteins, while unique and razor peptides were used for quantification of all other proteins. Downstream analysis was performed in R (v4.3.1) using limma (v3.56.2) ^51^ to calculate differentially abundant proteins. Moderated t-tests were applied with empirical Bayes shrinking of variance estimates (eBayes function) on a design matrix consisting of the contrasts of interest, corresponding to each compound vs DMSO, and degraders vs IAG933. Multiple hypothesis testing correction was performed by calculating Benjamini-Hochberg adjusted p-values for each contrast ^52^. The mean of the log transformed p values of the gene with the lowest p-adjusted value from the non-significant genes and log transformed p-values of the gene with the highest p-adjusted value from the significant genes was used to show the p-value threshold corresponding to an adjusted p-value of 0.05 on the volcano plots.

### RNA sequencing and bioinformatics analysis

Total RNA was purified from cells using Qiagen RNeasy plus kit according to the manufacturer’s instructions. Messenger RNA was purified from total RNA using poly-T oligo-attached magnetic beads. After fragmentation, the first strand cDNA was synthesized using random hexamer primers, followed by the second strand cDNA synthesis using dTTP to generate unstranded libraries. The library was checked with Qubit and real-time PCR for quantification and bioanalyzer for size distribution detection. Quantified libraries were pooled and sequenced on Illumina NovaSeq X Plus Series, with paired-end reads of 150nt (PE150).

Quality of paired-end reads was checked using FastQC v0.12.1. Quality and adapter trimming was performed using trim_galore v0.6.10 with default parameters. Pseudoalignment and transcript counting was performed on quality-trimmed fastq files using salmon v1.10.1 with parameters “-l A --validateMappings”. Further analysis was performed within R (v4.4.0). Gene-level counts were converted from salmon transcript counts using tximport ^53^ with options ignoreTxVersion=T, txOut=F, countsFromAbundance = “lengthScaledTPM”. Lowly-expressed genes that failed to meet the criteria > 10 counts in at least some samples and > 15 total counts were filtered out using filterByExpr from edgeR package v4.2.1 ^54^. Library sizes were normalised using calcNormFactors also from edgeR. Voom ^55^ was used to transform count data to log2 counts-per-million, estimate mean-variance relationship and compute observation-level precision weights. Limma ^51^ moderated t-test was subsequently applied with empirical Bayes shrinking of variance estimates to calculate adjusted p-values and logFC for each gene per contrast of interest, corresponding to each compound vs DMSO. Gene set enrichment analysis (GSEA) was performed using clusterProflier (version 4.16.0) ^56^ using the fast GSEA method ^57^. The full gene list for each contrast was ranked using the weighted metric - log10(pval) * sign(logFC) in descending order. Normalised Enrichment Score (NES) was calculated using the GSEA function from clusterProfiler with parameters eps=0, minGSSize=10, maxGSSize=500, with p-value adjustment method set to Benjamini-Hochberg and an adjusted p-value threshold of 0.05. Hallmarks gene signatures were obtained from MSigDb via the msigdbr package (version 25.1.1) with the addition of a custom YAP TAZ gene signature collated from the literature ^43,46–48^.

### Recombinant protein production and HTRF

His-tagged TEAD1 (209-426) was expressed in *E.coli* BL21(DE3) from pET28-His-TEV-G-TEAD1(209-426). Protein was purified by Ni^2+^-NTA affinity chromatography and purified by gel filtration in 50 mM Tris pH 7.5, 200 mM NaCl, 1 mM TCEP.

GST-tagged YAP1 (50-171) was expressed in *E.coli* BL21(DE3) Gold from pET28a-GST-GGS-3C-YAP1(50-171). The protein was purified by Glutathione Sepharose affinity chromatography, followed by gel filtration in 50 mM Tris, pH 8.0,150 mM NaCl, 5% Glycerol, 5 mM β-mercaptoethanol.

Biochemical IC_50_ values were determined by HTRF, using a TEAD1-YAP displacement assay. Briefly, 6nM TEAD1 His-TEV-G-TEAD1(209-426) was incubated with 6 nM YAP1 GST-GGS-3C-YAP1(50-171) plus DMSO or compound (11-doses, 10 uM highest concentration, 3-fold dilutions in technical duplicates) for 30 minutes at room temperature in OptiPlate 384-well Microplates (PerkinElmer, Cat.6007290). HTRF detection antibodies anti-His-Tb-Gold (Cisbio, 61HI2TLB) and anti-GST-d2 (Cisbio, 61GSTDLA) were added to the mixture and incubated for a further 2 hours at room temperature. Data was read on a PerkinElmer EnVision 2104 Multilabel Reader. IC_50_ values were calculated by nonlinear regression analysis using GraphPad Prism and at least two independent biological replicates were performed per compound.

### Statistical analyses and reproducibility

For immunofluorescence, cell viability and quantitative PCR assays data was completed with technical duplicates. Two independent biological replicates were completed, unless otherwise stated. For immunoblotting, the number of biological replicates is outlined in each figure legend.

## Supplementary Figures

**Supplementary Table 1:**
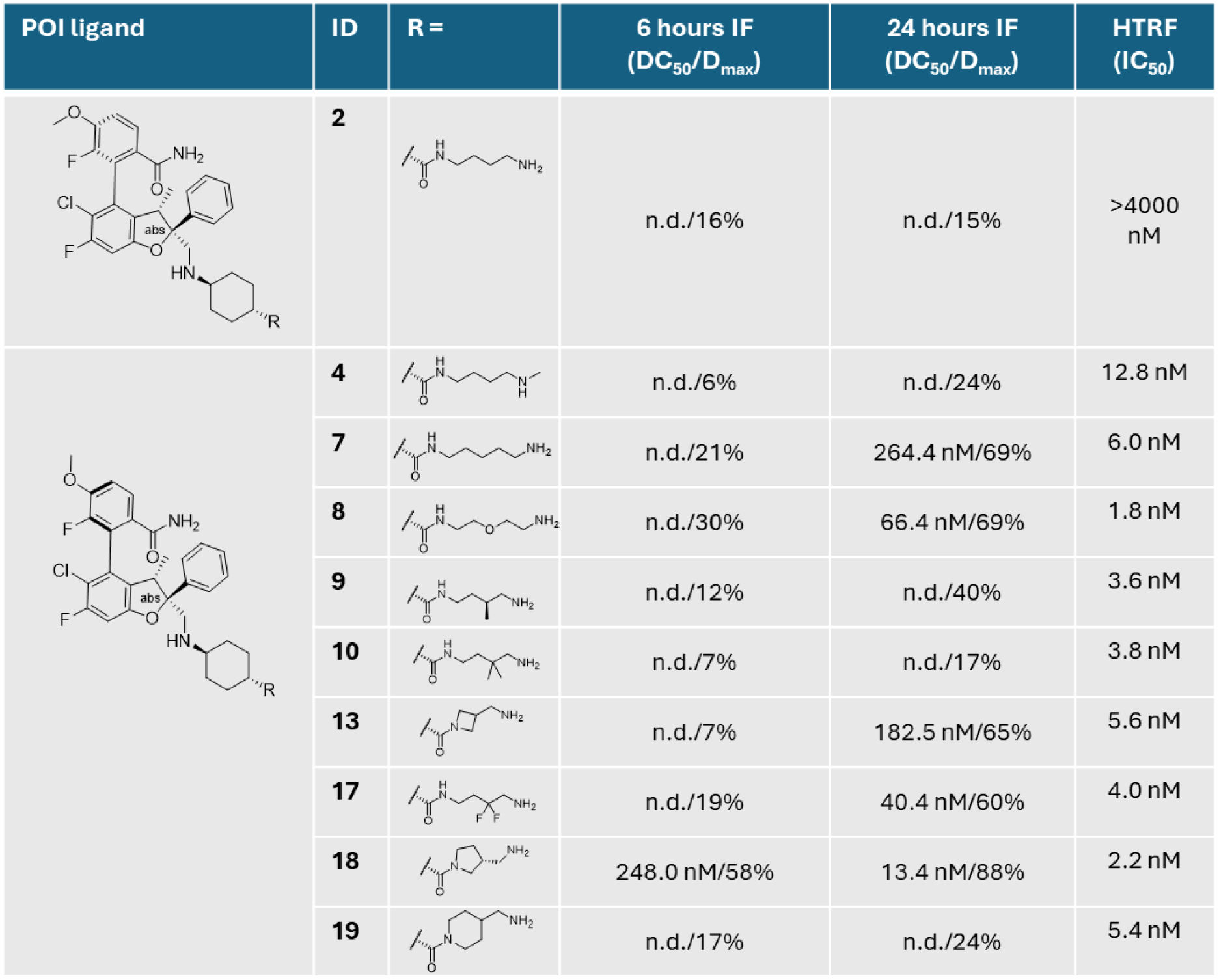
Data summary for TEAD degraders amine **2,** amine **4**, amine **7**, amine **8**, amine **G**, amine **10**, amine **13**, amine **17**, amine **18**, amine **1G**. Activity at 6 hours and 24 hours in NCI-H226 as determined by immunofluorescence; TEAD1 binding as measured via HTRF TEAD1-YAP displacement assay. IF data represents mean DC_50_ and D_max_ values for n = 1 or n = 2 biologically independent samples; HTRF data is mean IC_50_ values of n≥2 biological replicates. Graphs for amine **2** 24 hours immunofluorescence and HTRF are shown in Supp 3. Graph for amine **1** 24 hours immunofluorescence is shown in Fig 1a.

| POI ligand | ID | R = | 6 hours IF<br>(DC<sub>50</sub>/D<sub>max</sub>) | 24 hours IF<br>(DC<sub>50</sub>/D<sub>max</sub>) | HTRF<br>(IC<sub>50</sub>) |
| --- | --- | --- | --- | --- | --- |
|  | **2** |  | n.d./16% | n.d./15% | >4000 nM |
|  | **4** |  | n.d./6% | n.d./24% | 12.8 nM |
|  | **7** |  | n.d./21% | 264.4 nM/69% | 6.0 nM |
|  | **8** |  | n.d./30% | 66.4 nM/69% | 1.8 nM |
|  | **9** |  | n.d./12% | n.d./40% | 3.6 nM |
|  | **10** |  | n.d./7% | n.d./17% | 3.8 nM |
|  | **13** |  | n.d./7% | 182.5 nM/65% | 5.6 nM |
|  | **17** |  | n.d./19% | 40.4 nM/60% | 4.0 nM |
|  | **18** |  | 248.0 nM/58% | 13.4 nM/88% | 2.2 nM |
|  | **19** |  | n.d./17% | n.d./24% | 5.4 nM |

**Supplementary Figure 1:**
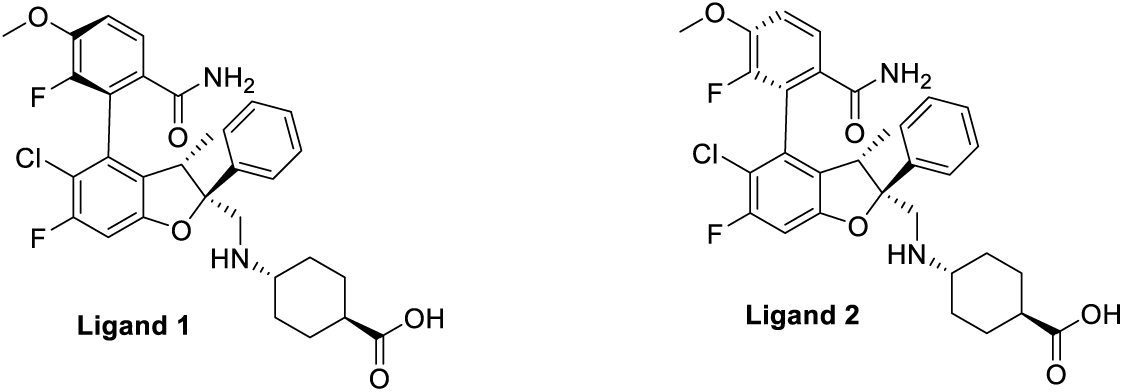
structure of POI ligand 1, and its non-TEAD binding atropisomer ligand 2

**Supplementary Figure 2:**
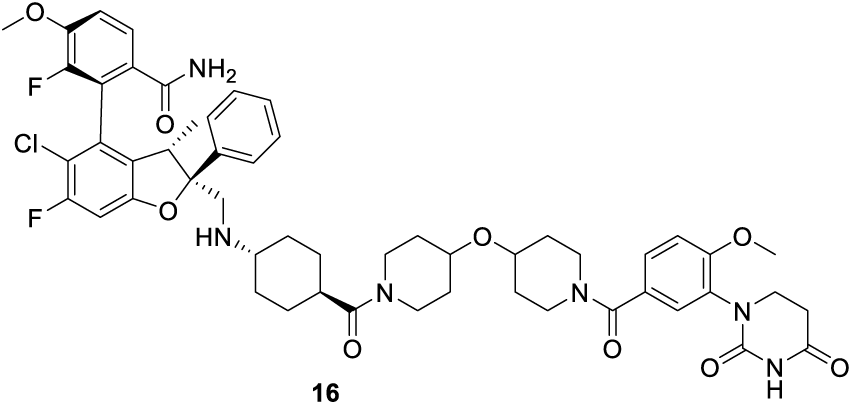
Structure of TEAD CRBN PROTAC **16**

**Supplementary Figure 3:**
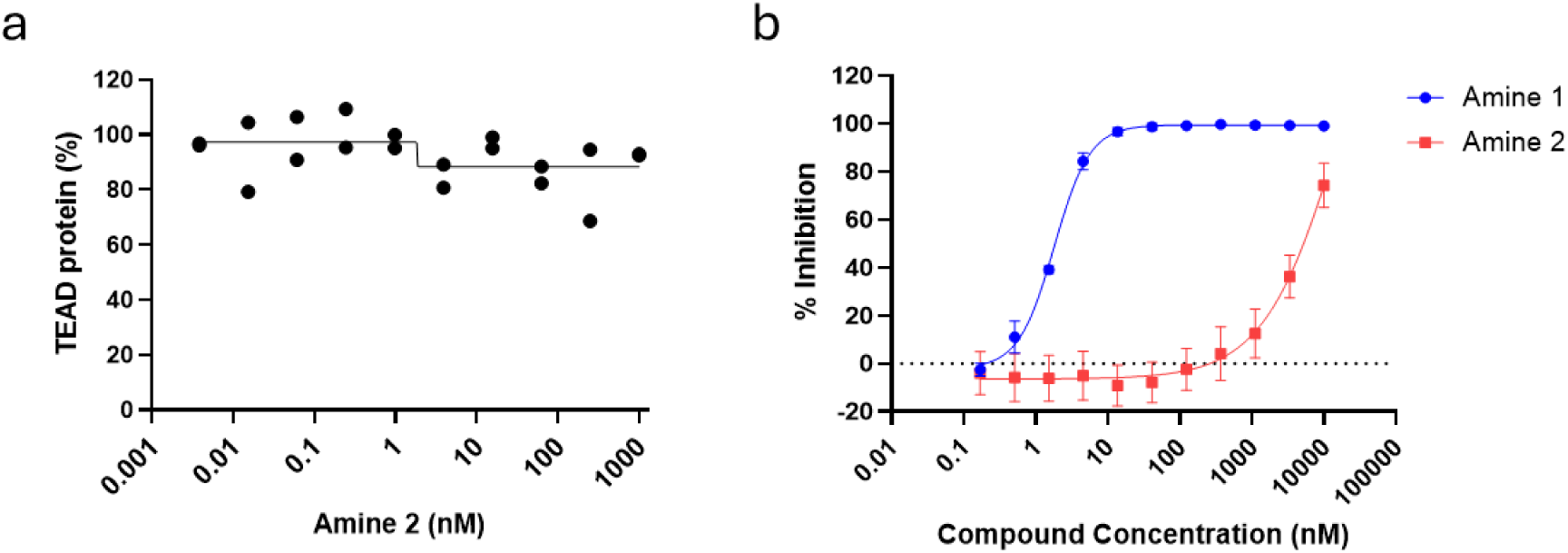
**a.** TEAD1 protein levels in NCI-H226, determined by immunofluorescence, following 24 hours treatment with a dose response of amine **2**. Data show the individual data points from n = 2 biologically independent samples. **b.** Biochemical HTRF TEAD1-YAP displacement assay following incubation with amine **1** and amine **2**. Data shown is the mean of four independent biological replicates ± SD. Resulting IC_50_ values are presented in Table 1 for amine **1** and Supp Table 1 for amine **2**.

**Supplementary Figure 4:**
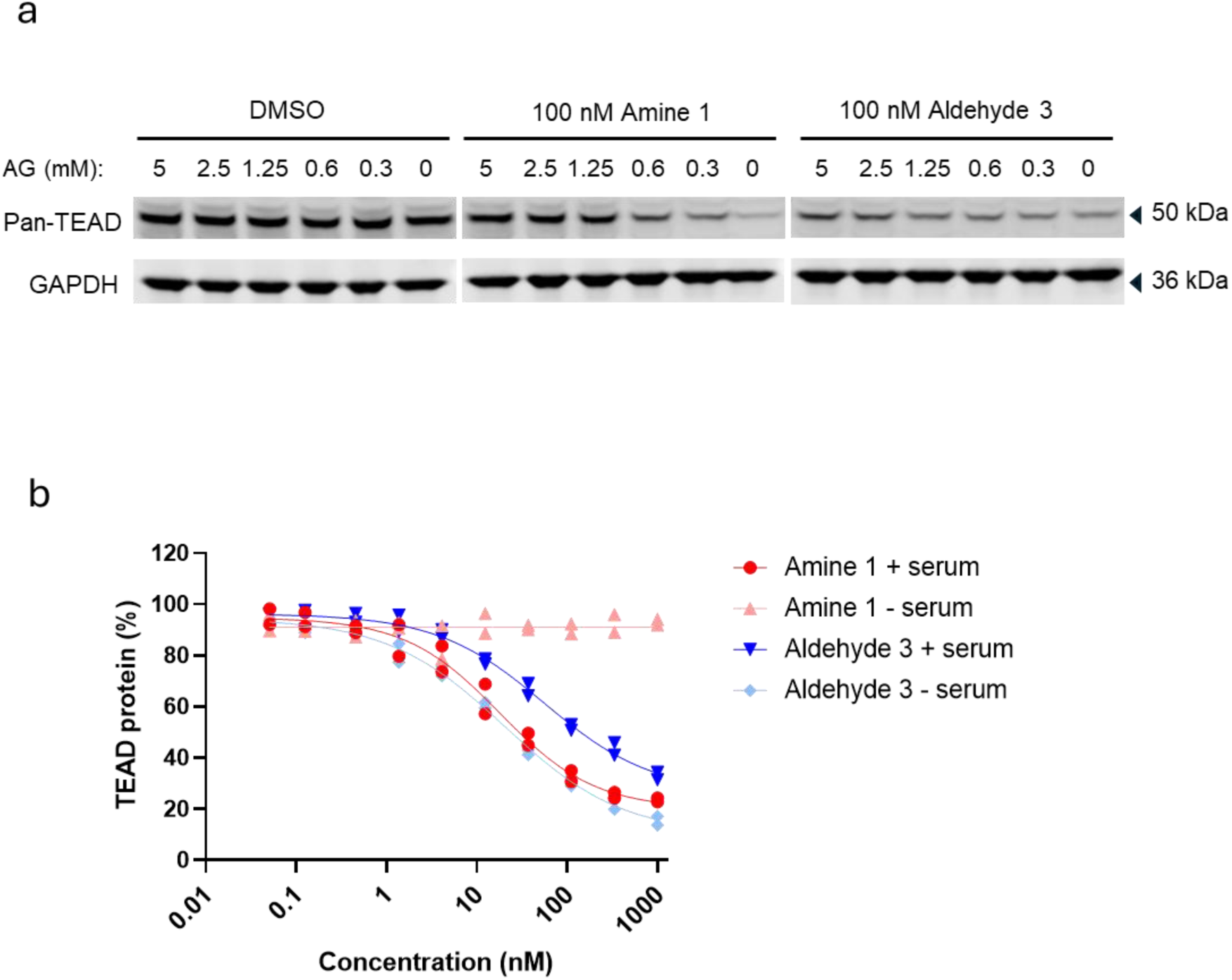
**a.** Representative immunoblot of pan-TEAD protein levels in NCI-H226 following 24 hours treatment with 100 nM amine **1** or aldehyde **3** in the presence of AG. Data associated with bar graph in Figure 2b. Data are from n = 2 biologically independent samples. **b.** TEAD1 protein levels in NCI-H226, determined by immunofluorescence, following 6 hours treatment with amine **1** and aldehyde **3** in standard or serum-free media. Data show the individual data points from n = 2 biologically independent samples.

**Supplementary Figure 5:**
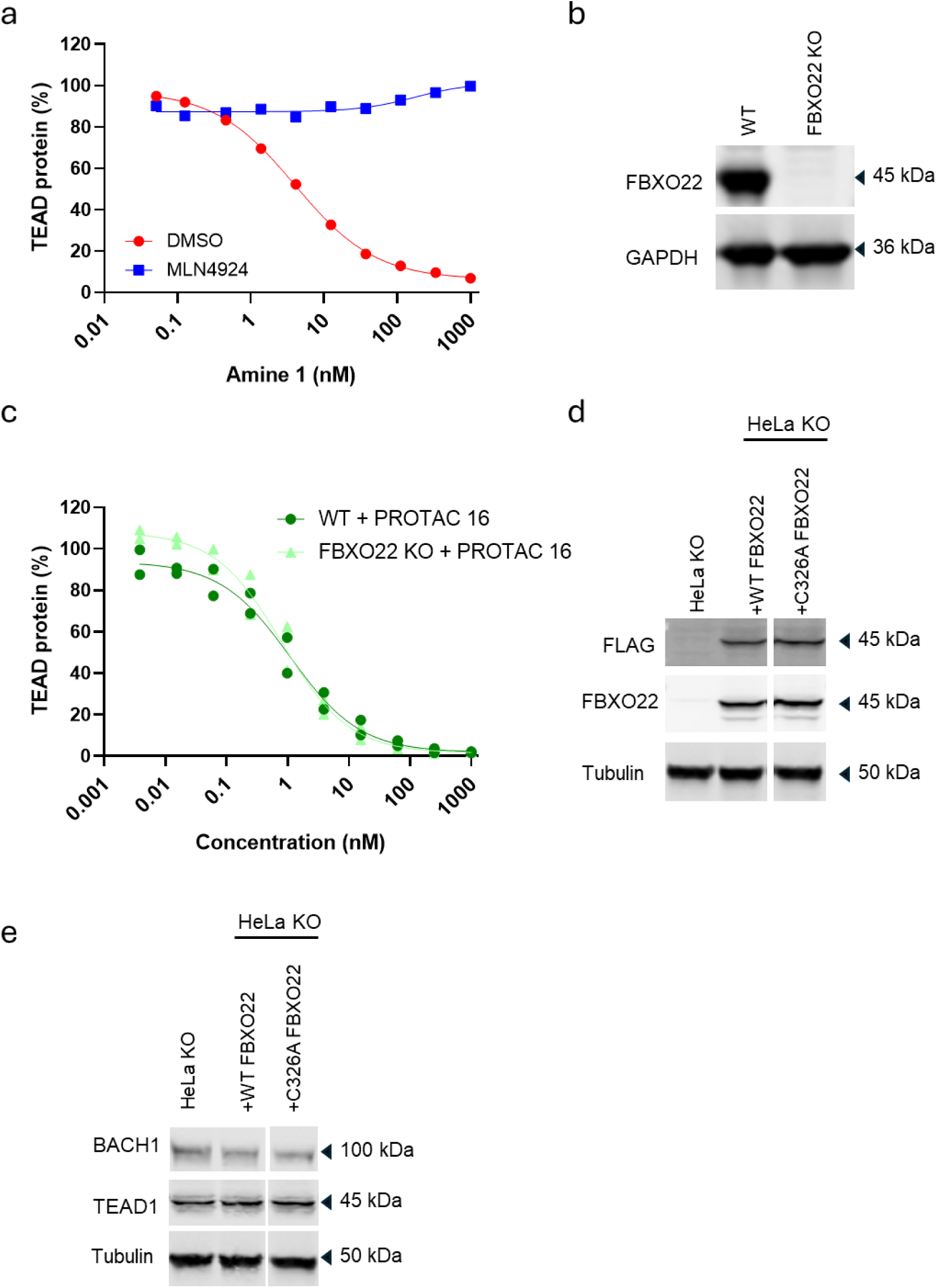
**a.** TEAD protein levels in NCI-H226, determined by immunofluorescence, following 1H pre-treatment with DMSO (red) or 3 µM MLN4924 (blue) then 24 hours treatment with amine **1**. Data are from n = 1 biologically independent samples. **b.** Immunoblot of FBXO22 protein levels in HeLa WT and FBXO22 KO cell lines. Data are from n = 1 biologically independent sample. **c.** TEAD protein levels in parental HeLa WT or FBXO22 KO cell lines, determined by immunofluorescence, following 24 hours treatment with a dose response of PROTAC **16**. Data show the individual data points from n = 2 biologically independent samples.

**Supplementary Figure 6:**
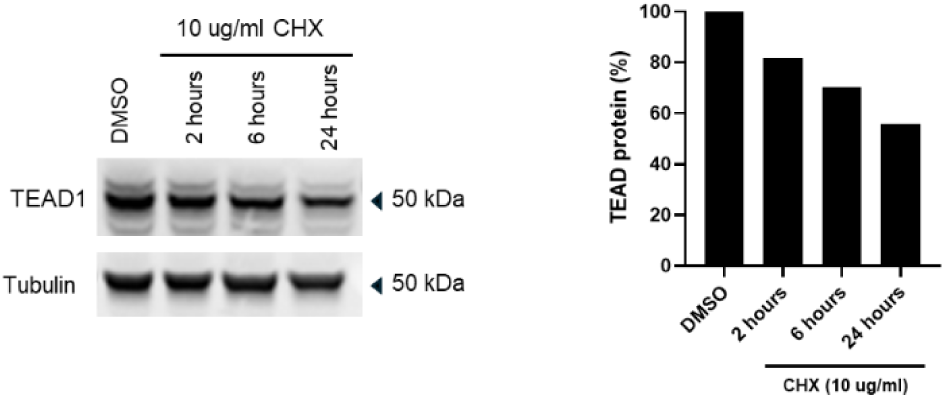
Immunoblot and quantification of TEAD protein levels in NCI-H226 following DMSO or 10 µg/ml cycloheximide over a time course. Data are from n = 1 biologically independent samples.

**Supplementary Figure 7:**
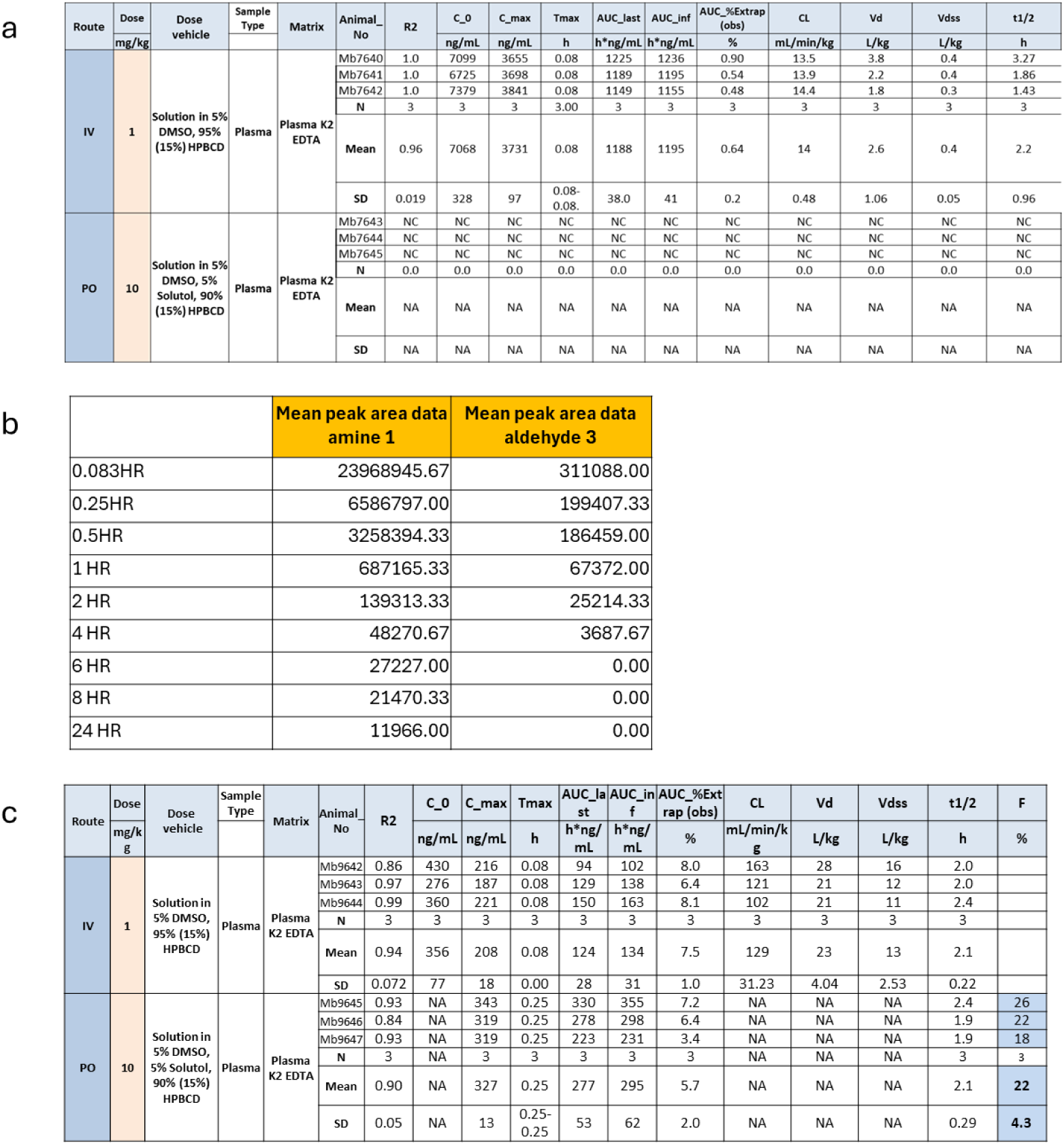
Pharmacokinetic (PK) data for amine **1,** its metabolite aldehyde **3**, and amine **17**. **a.** Details of PK experiment with amine 1 in CD-1 mice**. b.** Averaged peak area data for amine **1** and aldehyde metabolite **3** taken from mouse plasma during the amine **1** PK experiment **c.** details of PK experiment with amine **17** in CD-1 mice.

**Supplementary Figure 8:**
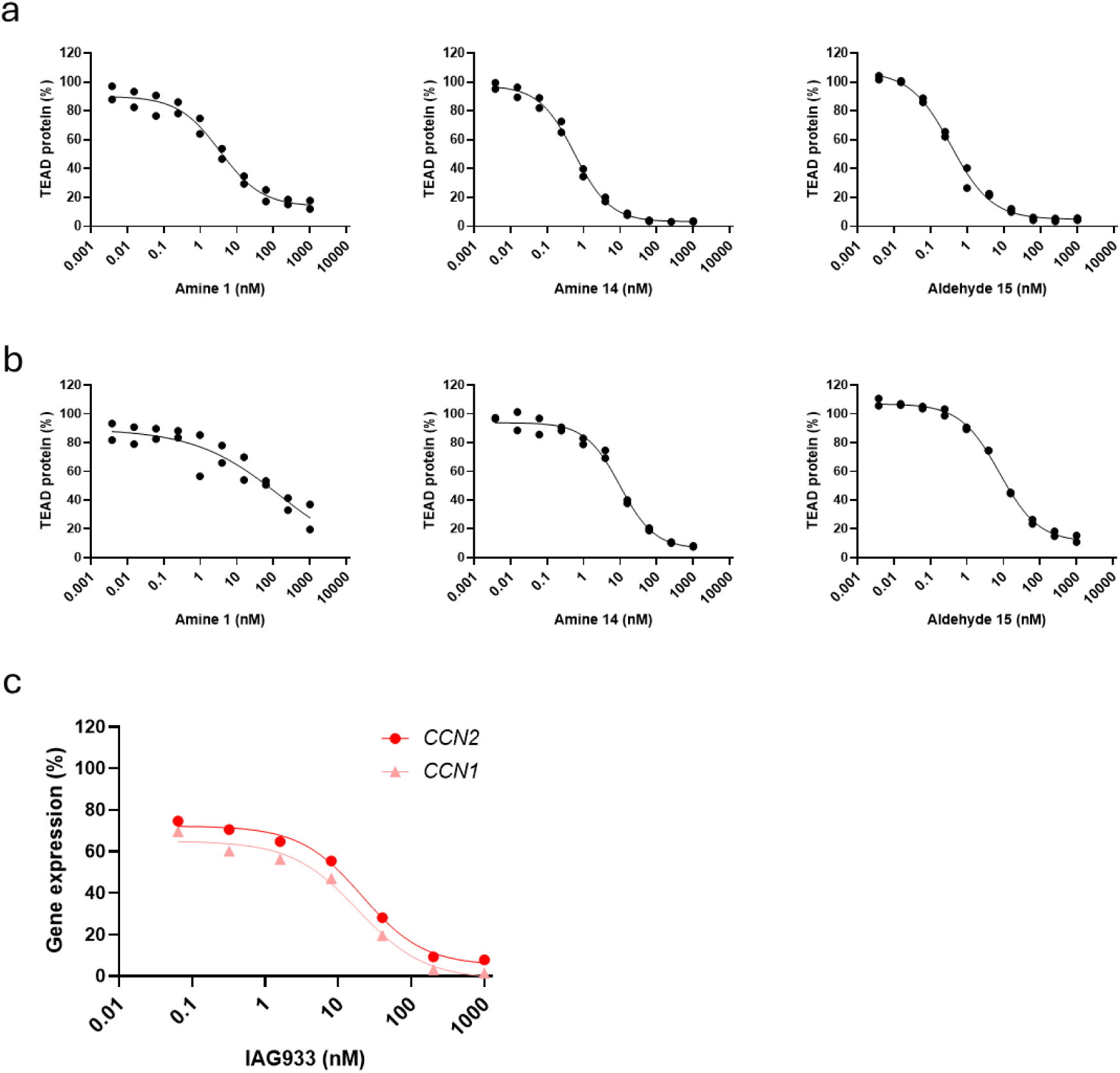
**a.** TEAD protein levels in NCI-H358, determined by immunofluorescence, following 24 hours treatment with amine **1** and **14** and aldehyde **15**. Data show the individual data points from n = 2 biologically independent samples. **b.** TEAD protein levels in PC-9, determined by immunofluorescence, following 24 hours treatment with a dose response of amine **1** and **14** and aldehyde **15**. Data show the individual data points from n = 2 biologically independent samples. **c.** *CCN1* and *CCN2* gene expression levels measured by qPCR in MSTO-211H following 24 hours treatment with a dose response of IAG933. Data show the individual data points from n = 1 biologically independent sample.

**Supplementary Figure 9:**
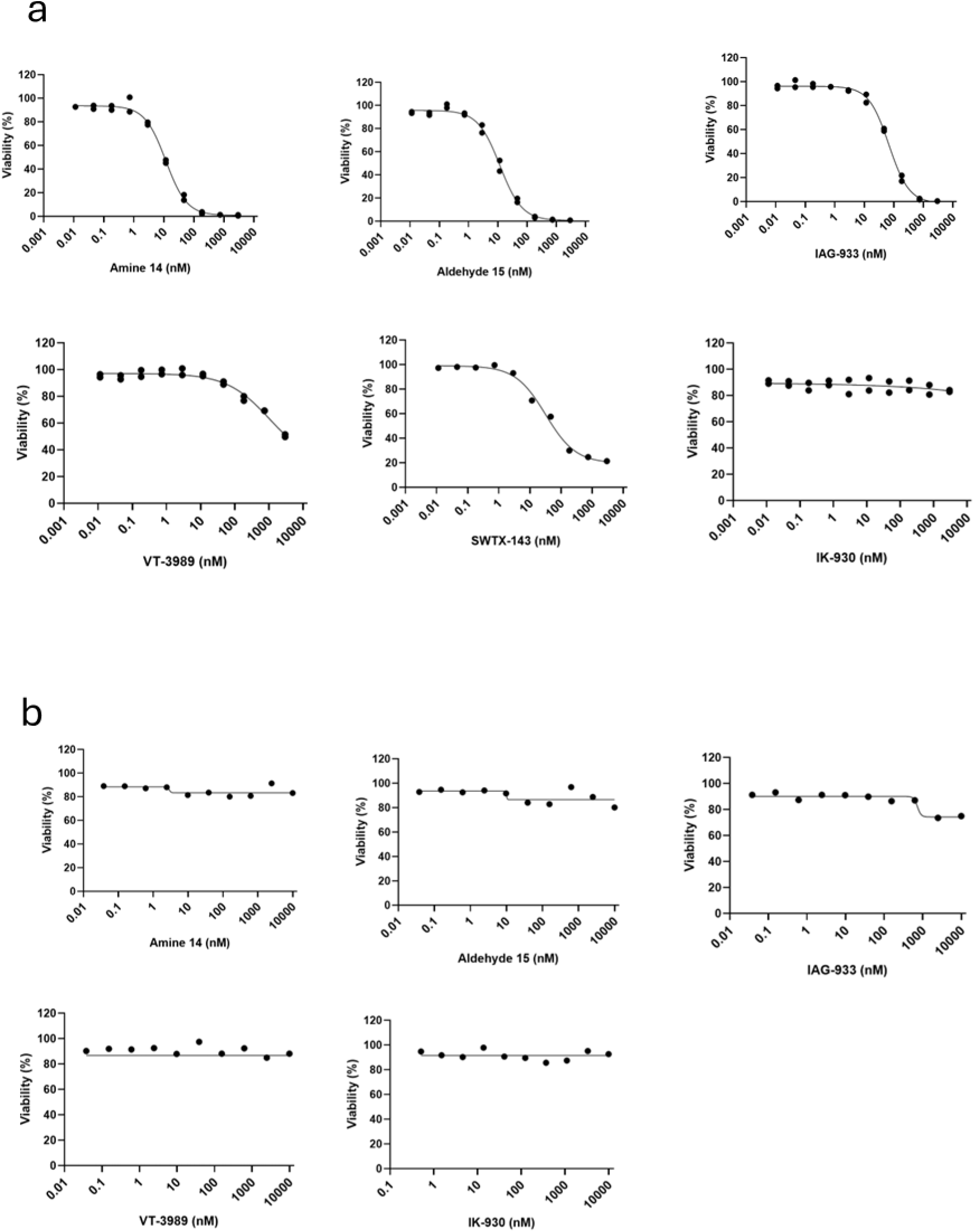
**a.** Cell viability in MSTO-211H, determined by CellTiter-Glo, following 120 h treatment with amine **14,** aldehyde **15,** IAG933, VT-3989, SWTX-143 and IK-930. Data show the individual data points from n≥1 biologically independent. **b.** Cell viability in HepG2, determined by CellTiter-Glo, following 120H treatment with amine **14,** aldehyde **15,** IAG933, VT-3989 and IK-930. Data are from n = 1 biologically independent sample.

**Supplementary Figure 10:**
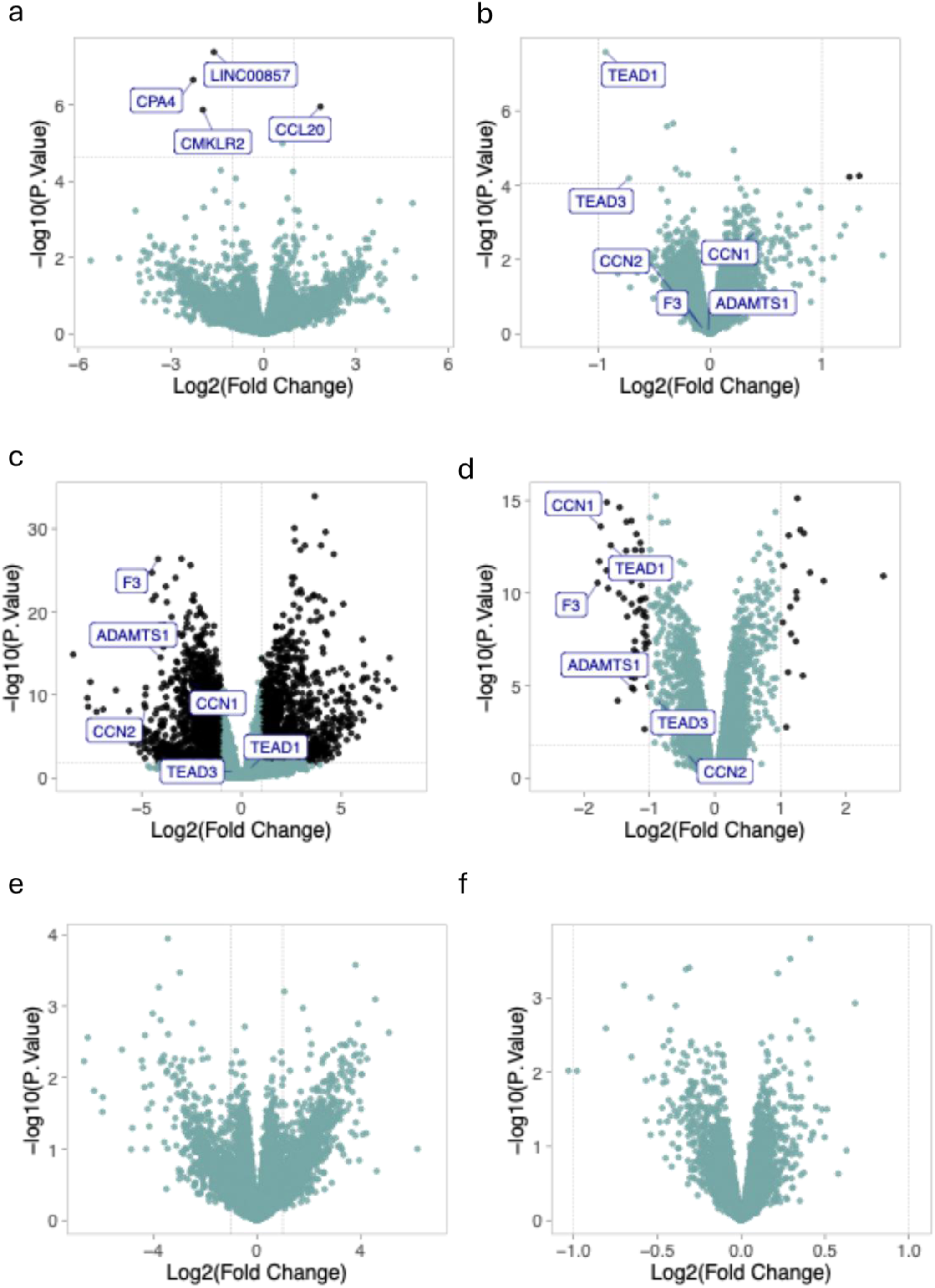
**a.** Volcano plot depicting RNA-seq comparison of amine 1 vs IAG933 after 24 hours treatment of NCI-H226 cells with 500 nM of each compound. Significantly changed genes were identified based on a threshold of adjusted p-value < 0.05 and absolute log_2_ fold change > 1. All significant genes are labelled in the plot. **b.** Volcano plot depicting proteomics comparison of 500 nM amine **1** vs 500 nM IAG933 after 24 hours treatment in NCI-H226 cells. Proteins were considered significantly altered if they had an adjusted p-value < 0.05 and absolute log_2_ fold change > 1. Hippo pathway-associated proteins TEAD1, TEAD3, CCN1, CCN2, F3, ADAMTS1 are labelled. **c.** Volcano plot depicting RNA-seq comparison of PROTAC **16** vs DMSO after 24 hours treatment of NCI-H226 cells with 500 nM of the compound. Significantly changed genes were identified based on a threshold of adjusted p-value < 0.05 and absolute log_2_ fold change > 1. Hippo pathway-associated proteins TEAD1, TEAD3, CCN1, CCN2, F3, ADAMTS1 are labelled. **d.** Volcano plot depicting proteomics comparison of PROTAC **16** vs DMSO after 24 hours treatment of NCI-H226 cells with 500 nM of the compound. Significantly changed proteins were identified based on a threshold of adjusted p-value < 0.05 and absolute log_2_ fold change > 1. Hippo pathway-associated proteins TEAD1, TEAD3, CCN1, CCN2, F3, ADAMTS1 are labelled. **e.** Volcano plot depicting RNA-seq comparison of amine **1** vs PROTAC **16** after 24 hours treatment of NCI-H226 cells with 500 nM of each compound. No significantly changed genes were identified based on a threshold of adjusted p-value < 0.05 and absolute log_2_ fold change > 1. **f.** Volcano plot depicting proteomics comparison of amine 1 vs PROTAC **16** after 24 hours treatment of NCI-H226 cells with 500 nM of each compound. No significantly changed proteins were identified based on a threshold of adjusted p-value < 0.05 and absolute log_2_ fold change > 1.

